# dsRADAR: Imaging and Quantifying Cellular dsRNA by Repurposing RNA Binding Proteins

**DOI:** 10.64898/2026.05.12.724404

**Authors:** Weina Cheng, Tyson D. Todd, Harshad Ingle, Angela M. Halstead, Megan T. Baldridge, José B. Sáenz, Jennifer M. Heemstra

## Abstract

Double-stranded RNA (dsRNA) is recognized by cellular receptors as a sign of viral infection, triggering the innate immune response. Increasing evidence shows that cellular dysregulation, for example in immune disorders and neurodegenerative diseases, can also lead to accumulation of endogenously produced dsRNA that stimulates a viral-like immune response. Additionally, dsRNA contamination in RNA therapeutics can lead to harmful side effects via a similar pathway. Despite the clinical relevance of dsRNA, reliable tools for its detection remain limited. At present, dsRNA detection relies almost exclusively on the monoclonal antibodies J2 and K1, which suffer from sequence bias and low sensitivity, limiting their reliability. To address this challenge, we aimed to repurpose naturally occurring dsRNA-binding domains (dsRBDs) to produce reliable, pan-specific affinity reagents for dsRNA. We first systematically screened the dsRBDs of the three human adenosine deaminases acting on RNA (ADARs). This analysis identified ADAR3 dsRBDs as promising candidates due to their reduced sequence dependence compared to the dsRBDs of ADAR1 and ADAR2. We then engineered ADAR3-derived dsRBD constructs having varying linker lengths and domain combinations, allowing us to specifically vary the length cutoff of dsRNA detected, thus creating dsRNA accumulation detected by ADAR3 RBDs (dsRADAR) affinity reagents. Finally, we demonstrate the superior performance of dsRADAR over currently available dsRNA antibodies in a cell model of viral infection and a tissue model of gastric inflammation. Together, dsRADAR provides a sensitive and reliable approach for imaging and quantifying diverse dsRNA structures in a variety of biological contexts.

**Graphic Abstract:** 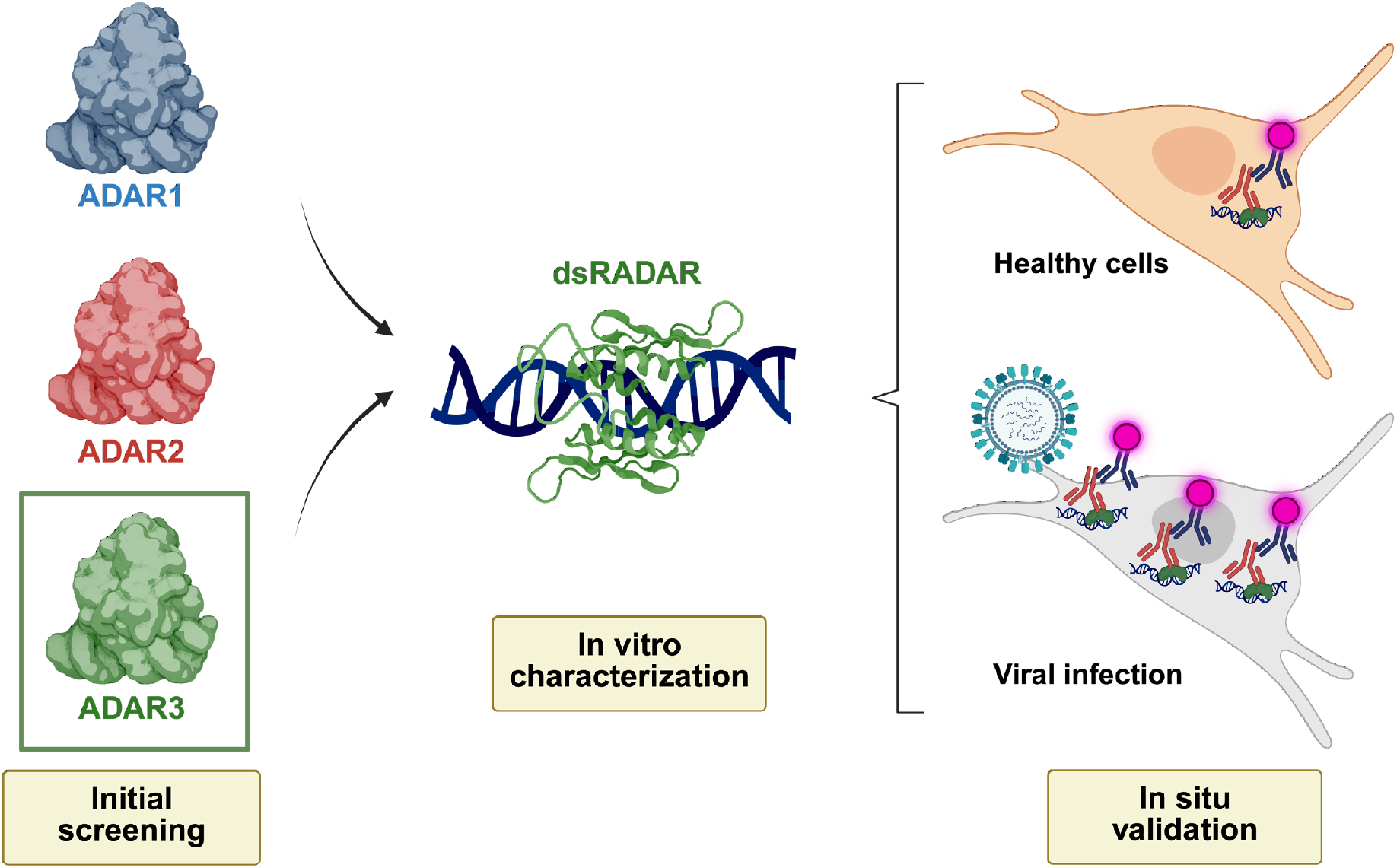

## Introduction

Cellular double-stranded RNA (dsRNA) accumulation has historically been associated with viral infection and can arise from multiple types of viruses. In the case of dsRNA viruses, such as reoviruses, replication and transcription occur primarily within the viral core to prevent exposure of dsRNA and subsequent immune activation.^1^ Thus, in these infections, recognition by cellular dsRNA sensors happens predominantly through the leakage of viral dsRNA from the capsid.^1^ Both positive- and negative-strand ssRNA viruses produce dsRNA as replicative intermediates that can be recognized by cellular dsRNA sensors, such as MDA5 and RIG-I.^2–5^ In the case of DNA viruses, such as adenovirus, a significant level of dsRNA has been shown to accumulate through bidirectional transcription, and this RNA is recognized by RIG-I.^6–8^ Upon binding to viral dsRNA, these cellular sensors activate various pathways of the innate cellular immune response, including antiviral and inflammatory signaling, inhibition of cell growth, or even cell death to prevent viral replication.^3,4,9–11^ Therefore, dsRNA detection serves as a reliable indicator of viral infection for both research and diagnostic purposes.

While dsRNA is primarily associated with viral infections, recent studies have shown that endogenous dsRNA accumulation can also result from cellular dysregulation and thus may serve as a biomarker of disease. In these cases, dsRNA originates mostly from non-coding regions of the transcriptome, such as short and long interspersed elements (SINEs, LINEs), endogenous retroviruses (ERVs), and Alu elements.^12^ Once transcribed, cellular RNAs can undergo adenosine-to-inosine (A-to-I) editing, which functionally recodes adenosine to behave similarly to guanosine and serves to disrupt the structure of dsRNA and lower its immunogenicity.^13^ The accumulation of endogenous dsRNA or dsRNA-like species due to dysregulation of cellular processes can activate immune responses analogous to those triggered by exogenous viral dsRNA.^12,14,15^ This cellular dysfunction can be traced to multiple pathways, including dysregulated epigenetic control, inhibition of proper splicing, and defects in RNA modifications or proper RNA degradation.^16,17^ Such aberrant immune activation in the absence of viral infection often leads to the pathogenesis of immune disorders, such as Aicardi-Goutières syndrome. Additionally, an increasing number of studies suggest a close association between dsRNA dysregulation and various diseases, including neurodegenerative disorders like Alzheimer’s disease and epithelial cancer, particularly pancreatic cancer.^18–20^

Quantifying dsRNA is also important for detecting contamination generated during *in vitro* transcription (IVT) to produce RNA vaccines and therapeutics. For instance, T7 RNA polymerase can initiate at the ends of DNA templates in a promoter-independent manner, producing antisense RNA that anneals with sense RNA to produce full-length dsRNA.^21^ Additionally, once ssRNA is transcribed, the polymerase can undergo template switching to the nascent RNA strand if regions of complementary RNA sequence are present between the end of the transcript and some internal site within the transcript, producing long dsRNA though self-templated extension.^21–24^ Similar to viral dsRNA, these transcriptional byproducts can stimulate the immune system and cause inflammation and other negative side effects.^22^

Together, the role of dsRNA in infection, disease, and therapeutics creates a critical need for efficient methods to detect the presence of dsRNA both *in vitro* and in cells and tissues. However, despite their importance, current detection methods remain limited. For *in vitro* dsRNA detection, the most commonly used technique is RNA extraction. However, studying dsRNA using this method is notoriously challenging as dsRNA secondary structure is sensitive to environmental conditions. The processes involved in isolating and handling cellular material may inadvertently modify, degrade, or reassemble dsRNA, making it difficult to accurately identify and quantify native dsRNA that existed in the cellular context compared with artifacts produced during processing.^25,26^ Additionally, extraction erases the important insights that could be gained by observing dsRNA subcellular localization and cell-to-cell variation, which may be especially relevant in tissues which are typically composed of several different cell types. In an attempt to address these limitations and enable *in situ* detection, monoclonal antibodies (mAb) J2 and K1 were developed and first used in 1991 by Schönborn et al., representing the earliest attempt to visualize dsRNA in fixed cells.^27^ While antibody-based detection *in situ* preserves the localization and structure of dsRNA in the cellular environment, subsequent studies have shown that these antibodies suffer from sequence specificity and low sensitivity. In particular, mAb J2 preferentially binds AU-rich dsRNA, recognizing a specific sequence pattern, A2-N9-A3-N9-A2, consisting of two A dyads flanking an A triad.^28^ Furthermore, J2 loses binding activity completely when GC content exceeds 63% and exhibits reduced affinity for dsRNA motifs shorter than 30-40 bp. This renders several classes of naturally occurring and biologically important dsRNAs undetectable by J2.^27,29^ The other mAb, K1, shows a preference for poly(I)·poly(C), which was the substrate used to derive this antibody but does not exist naturally in cells.^27–29^ Together, immunoassays using J2 and K1 still face various challenges and limitations, creating an ongoing need for improved methods for recognizing and quantifying dsRNA *in vitro* and in cells and tissues.

To address this need, we sought to develop a novel method for dsRNA detection by repurposing naturally evolved dsRNA binding domains (dsRBDs). We were particularly drawn to the dsRBDs of the enzyme family of adenosine deaminases acting on RNA (ADARs). ADARs are evolutionarily conserved RNA-editing proteins that convert adenosine to inosine within double-stranded RNA regions. As part of this function, their dsRBDs innately exhibit strong and specific affinity for dsRNA duplex regions, and have been shown to function independently as dsRNA binders without the catalytic deaminase domain.^30–33^ We initiated our study by screening all dsRBDs from the three human ADAR homologs and found that ADAR3 dsRBDs have the broadest sequence tolerance, thus serving as an ideal starting point for repurposing into a novel anti-dsRNA “antibody.” Based on this promising initial result, we were able to engineer and biophysically characterize dsRNA Accumulation Detected by ADAR RBDs (dsRADAR), two novel affinity reagents derived from the dsRBDs of ADAR3. dsRADARs show robust binding affinity and high specificity toward dsRNA over other nucleic acids, minimal sequence bias, and tunable dsRNA length detection cutoffs of 8bp and 25bp through variation of the linker between the two dsRBDs. We then developed an immunostaining workflow to interrogate dsRNA accumulation in virus-infected cell lines and diseased tissue samples to validate performance of the affinity reagents in biologically relevant contexts. dsRADAR consistently detected dsRNA accumulation in varied contexts and showed higher sensitivity than either the J2- or K1-based approaches. Together, these findings establish dsRADARs as a sensitive, sequence-independent, and biologically compatible approach for visualizing and quantifying dsRNA *in vitro and in situ*. This provides a powerful alternative to existing antibody-based dsRNA detection methods and creates a broadly applicable tool for studying dsRNA accumulation related to natural cellular processes, viral infection, and disease, as well as for detecting dsRNA contamination in RNA therapeutics.

## Results

### 1. Screening and engineering initial dsRBD constructs

The ADAR family provides a group of naturally occurring dsRNA binding domains that were selected as an initial starting point for developing a sequence-independent dsRNA recognition tool. Every human ADAR contains two or more dsRBDs that recognize A-form dsRNA duplexes with conserved structural motifs, but each ADAR exhibits different catalytic activities and RNA substrate preferences.^30^ ADAR1 and ADAR2 are catalytically active and have been shown to display context-dependent editing specificities.^34^ Previous studies have demonstrated that the deaminase domain primarily determines the editing specificity, while the dsRBDs contribute to recognition by dynamically scanning along the dsRNA structure until the catalytic domain finds its target site.^35–38^ This model suggests that the sequence preference is primarily dominated by the deaminase domain, which makes the isolated dsRBDs promising candidates toward our goal of creating universal dsRNA affinity reagents. In contrast to ADAR1 and ADAR2, ADAR3 is catalytically inactive but retains dsRNA binding ability, suggesting that its dsRBDs might also be good candidates for generating a dsRNA-binding affinity reagent.^39^ Therefore, we initiated a comparative screen on isolated ADAR dsRBD constructs using microscale thermophoresis (MST) to identify those with desired binding properties (Fig. 1, Fig. 2b-c).

**Figure 1.**
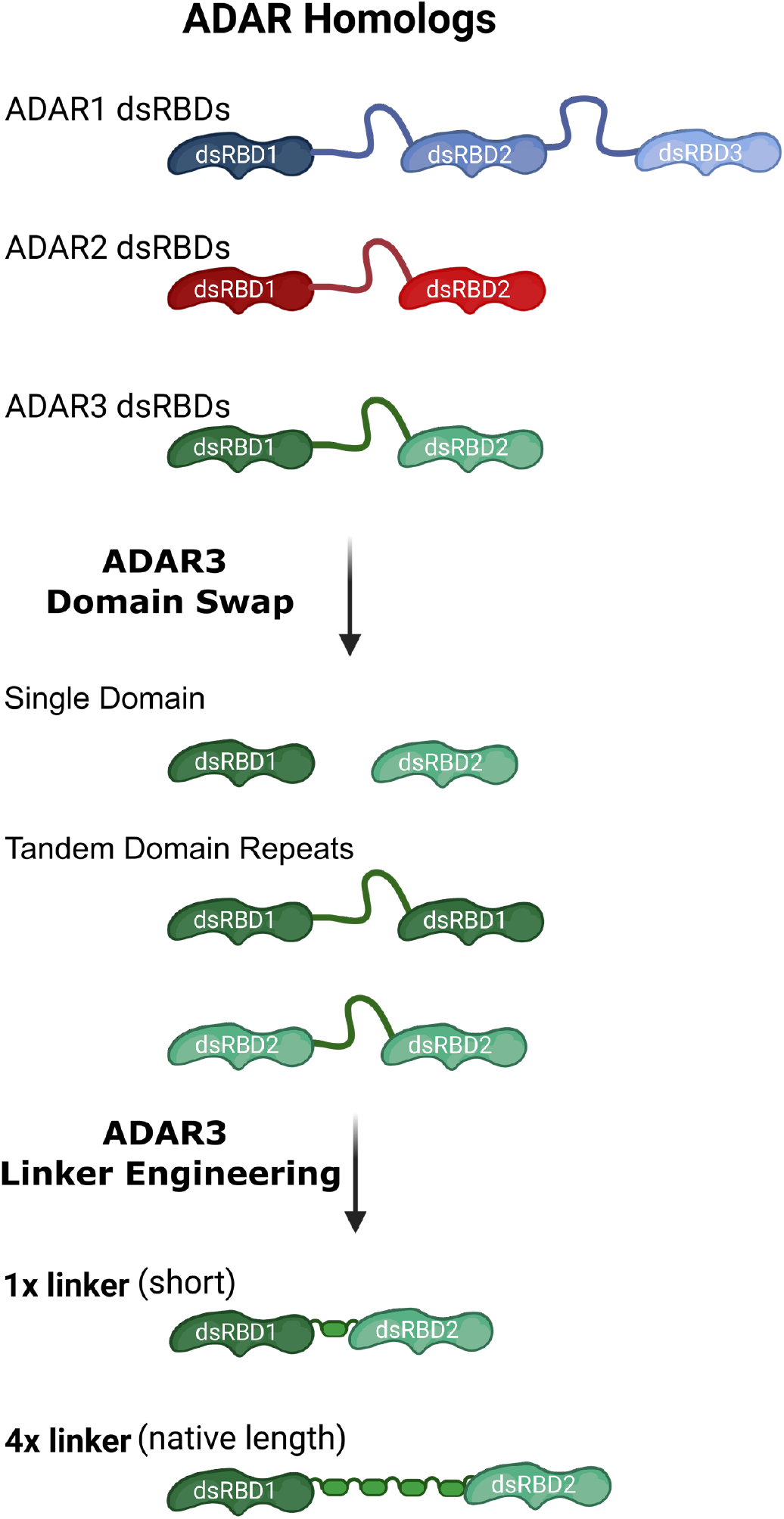
Engineering of ADAR dsRBD constructs for dsRNA binding. dsRNA-binding domains (dsRBDs) from human ADAR1, ADAR2, and ADAR3 were expressed and tested for binding with two randomized sequence dsRNA hairpins (rsRNA-1 and 2). Microscale thermophoresis (MST) analysis identified ADAR3 dsRBDs as having the highest sequence tolerance. ADAR3 variants, including single domains and tandem domain repeats, were then tested to assess interdomain cooperativity. ADAR3 variants having 1x and 4x repeats of a synthetic linker were constructed for further screening.

**Figure 2.**
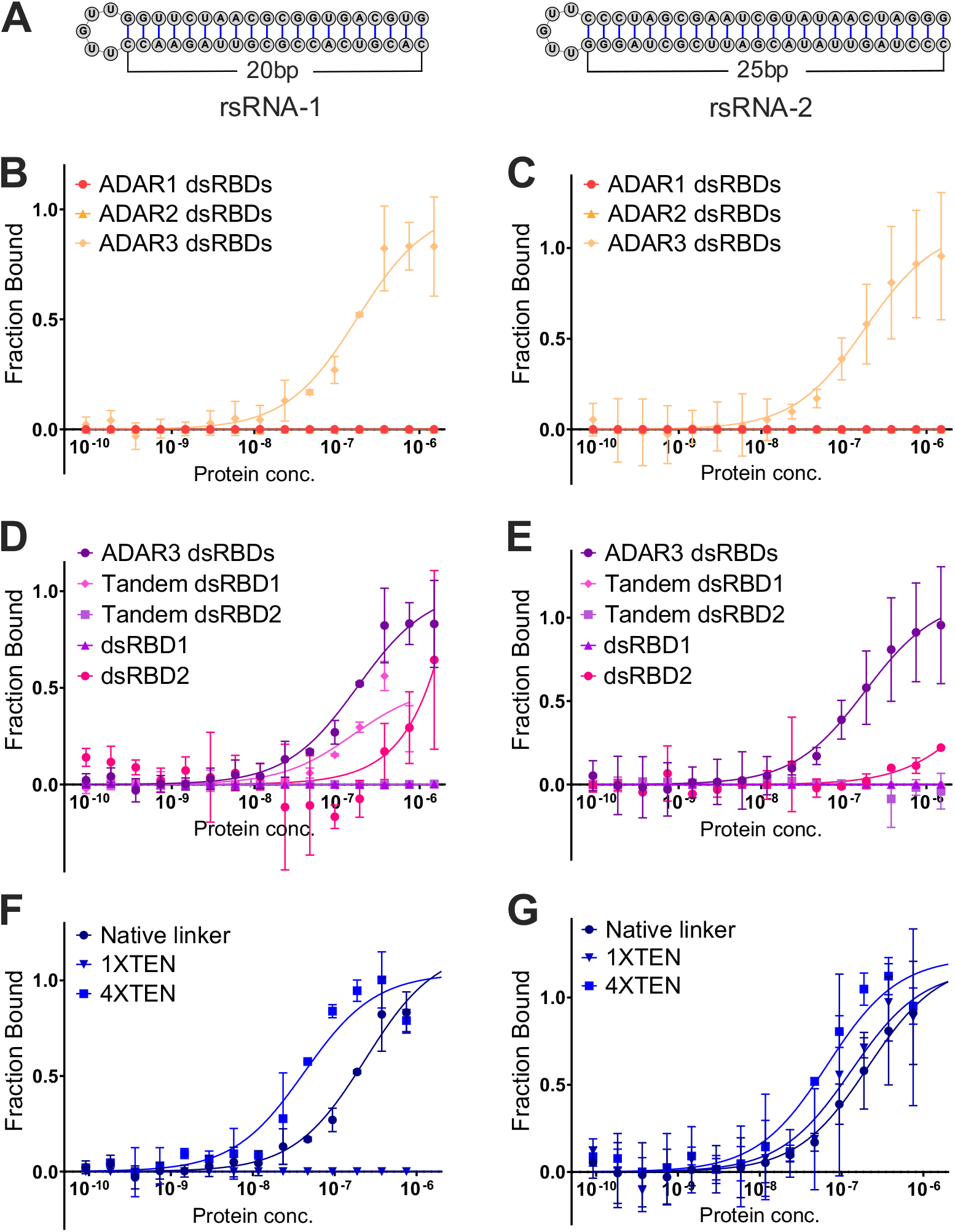
Characterization of dsRBD binding to random sequence RNAs. (a) Sequence and secondary structure of rsRNA-1 and rsRNA-2. Both RNA substrates contain a 3’ fluorescent label. (b,c) MST binding curves for dsRBDs of ADAR1, ADAR2, and ADAR3 measure with the (b) randomized dsRNA substrate rsRNA-1 and (c) the randomized dsRNA substrate rsRNA-2. (d,e) MST binding curves for ADAR3 variants, including the wild type dsRBDs, tandem dsRBD1, tandem dsRBD2, isolated dsRBD1, and isolated dsRBD2 measured with (d) rsRNA-1 and (e) rsRNA-2. (f,g) MST binding curves of dsRADAR variants having native, 1x, and 4x linkers measured with (f) rsRNA-1 and (g) rsRNA-2. All MST experiments were performed in 20 mM Tris, pH=7.6, 125 mM NaCl, 30 mM KCl, 5 mM MgCl_2_, 0.5 mM DTT.

Two independently randomized dsRNA hairpins, rsRNA-1, containing a 25bp duplex region with 56% GC content, and rsRNA-2, containing a 20bp duplex region with 47% GC content, were used to conduct this initial screening (Fig. 2a). Because each hairpin contains a different, non-overlapping randomized sequence, the possibility that both RNAs would coincidentally match a homolog’s intrinsic sequence preference is slim. Therefore, we reasoned that an ADAR dsRBD that binds both randomized substrates with comparable affinity is unlikely to be recognizing a specific sequence and is instead more likely to bind dsRNA in a sequence-independent manner. We expressed recombinant dsRBD fragments from all three ADARs in *E. coli* and measured their binding affinities to both rsRNA-1 and rsRNA-2 (Fig. 1, Fig. 2b-c). Among the three homologs, only the dsRBDs from ADAR3 exhibited strong binding to both target substrates, with *K*d values of 150 nM and 170 nM for rsRNA-1 and rsRNA-2, respectively (Fig. 2b-c, Supplementary Table S2). This observation suggests that, although the sequence preference of ADAR1 and ADAR2 is presumed to be determined predominantly by the deaminase domain, their dsRBDs may still contribute to RNA substrate selection. Fortuitously, ADAR3 dsRBDs do not seem to have this same sequence bias in our assay system and thus they were selected for further development.

We next investigated whether these domains function cooperatively or independently by purifying each individual dsRBD and assaying for binding with the same dsRNA substrates (Fig. 2d-e). The isolated dsRBD1 and dsRBD2 of ADAR3 appear to have negligible binding affinity to the rsRNA-2, and only dsRBD1 retained weak affinity to rsRNA-1, suggesting that cooperative interdomain and/or multivalent interactions are required for robust dsRNA binding by these domains (Fig. 2d-e). Tandem dsRBD repeat constructs of ADAR3 dsRBD1 and ADAR3 dsRBD2 were also tested to compare with the native dsRBD region containing both domains (ADAR3 dsRBD1-dsRBD2). Only the native dsRBD construct containing both dsRBD1 and dsRBD2 exhibited high-affinity binding to both RNA substrates (Fig. 2d-e). Even though the dsRBD1 tandem repeat construct retained its interaction with rsRNA-1, neither tandem repeat construct showed interactions with rsRNA-2. Together, these observations indicate that ADAR3 dsRBD1 and dsRBD2 act distinctly but also cooperatively to interact with dsRNA and maintain a broad RNA substrate scope.

### 2. Tunable dsRNA length binding of dsRADAR through linker optimization

With the ideal combination of dsRBDs identified, we next turned to exploring the linker between these constructs to test for length and composition dependence of the RNA binding interaction. We replaced the native linker with synthetic flexible linkers of varying lengths (1x and 4x repeats of a 20 amino acid sequence), where the 4x linker approximates the natural 82 amino acid linker length in ADAR3, and 1x represents the shortest reasonable linker unit between these two domains. While constructs containing the native linker or synthetic linkers exhibited comparable binding affinities by MST, those containing synthetic linkers consistently showed improved solubility and yield from recombinant expression. We noted that only the 4x variant maintained robust, near-native dsRNA-binding affinity to both rsRNA-1 and -2 with *K*d values of 54nM ± 20nM and 67nM ± 30nM, respectively (Fig. 2f-g, Supplementary Table. S2), whereas the 1x construct showed diminished binding to rsRNA-1. We reasoned that, as rsRNA-1 contains a shorter duplex region, shortening the linker may restrict the relative orientation and positioning of the dsRBDs on the duplex and prevent the 1x construct from forming favorable cooperative contact with rsRNA-1. In contrast, rsRNA-2, containing a slightly longer duplex, is permissive to a more optimal binding orientation for the 1x construct while both rsRNAs facilitate optimal binding for the 4x construct.

Based upon these observations, we hypothesized that two binding modes between dsRADAR and dsRNA are possible (Fig. 3c). The first is clamp mode, in which the two dsRBDs interact with opposite faces of the same region of dsRNA, effectively “clamping” the duplex between the two dsRBDs. Alternatively, in series mode, the two dsRBDs can bind adjacent segments of the same dsRNA along the helical axis, which extends the binding footprint. Structural studies of ADAR1 and ADAR2 dsRBDs indicate that a single dsRBD typically contacts around 8-10 bp of RNA.^30,35^ Thus, we hypothesize that our longer synthetic linker (4x) provides sufficient flexibility for the ADAR3 dsRBDs to bind in either clamp mode or in series mode, whereas the shorter (1x) linker prevents optimal positioning and orientation for clamp mode and makes series mode the preferred conformation.

**Figure 3.**
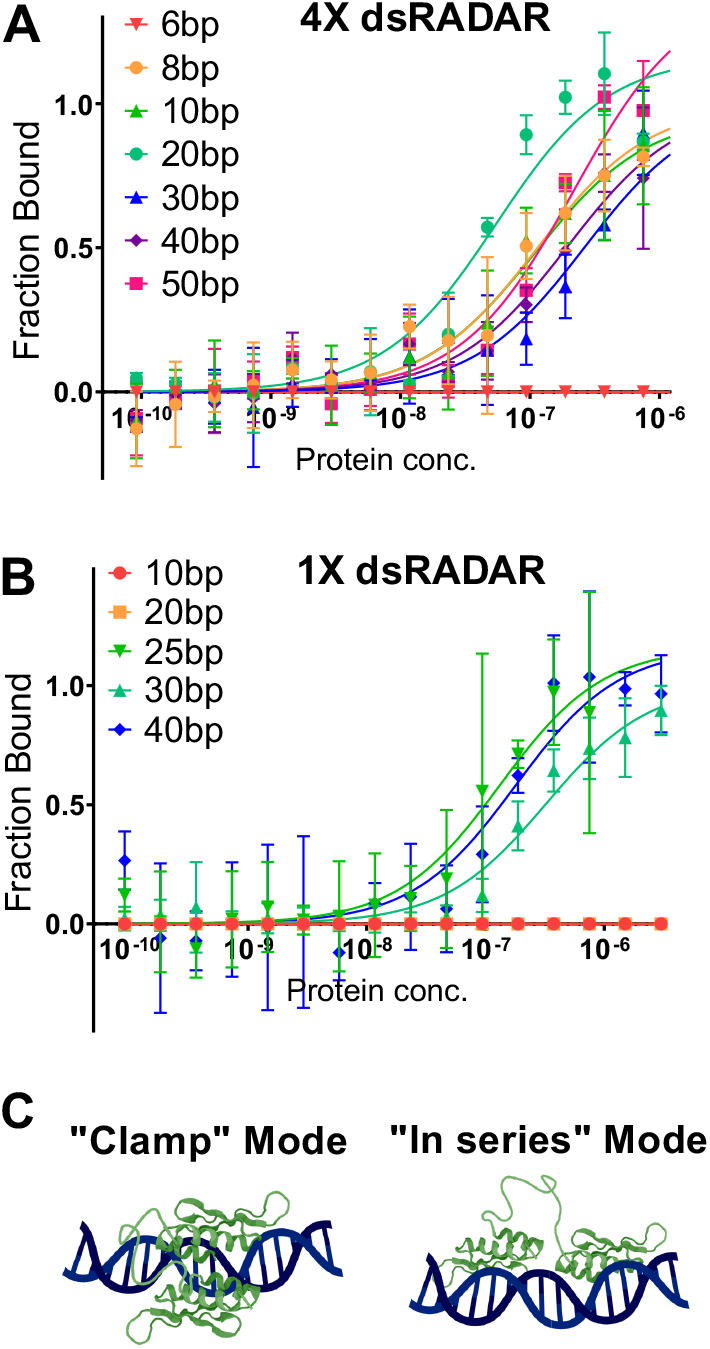
Tunable dsRNA length requirement of dsRADARs through linker optimization. MST binding curves of RNAs having varying duplex length with (a) 4x dsRADAR and (b) 1x dsRADAR. (c) Proposed models for two possible dsRADAR-dsRNA binding modes. All MST experiments were performed in 20 mM Tris, pH 7.6, 125 mM NaCl, 30 mM KCl, 5 mM MgCl_2_, 0.5 mM DTT.

To experimentally test these models further and define the minimal dsRNA length required for stable binding, we synthesized a panel of dsRNA substrates containing duplex regions ranging from 6 to 50 bp (Fig. S1a). We quantified binding affinity using MST and observed that the 4x construct retains robust binding with comparable *Kd* values across substrates having duplex lengths ranging from 8 to 50 bp (Fig. 3a, Supplementary Table. S1). This result agrees with our structural prediction that the 4x construct requires a minimum duplex length of 8-10 bp to form a stable “clamp” mode binding complex. In contrast, the shortened 1x variant only showed binding to duplexes of 25 bp or longer, nicely aligning with the results of our initial screening and our proposed “series” binding mode (Fig. 3b).

### 3. dsRADAR binds dsRNA in a nearly sequence agnostic manner

Virus-derived and endogenous dsRNAs exhibit a diverse range of sequence composition, GC content, base-pairing stability, and editing hotspots. Thus, an effective affinity reagent for dsRNA should have minimal sequence bias, which is unfortunately not the case for the currently available antibodies. We were encouraged by the ability of the ADAR3 dsRBDs to bind both initial screening RNA substrates (rsRNA-1 and -2) and variable length substrates with high affinity, and sought to more fully characterize the sequence tolerance of dsRADAR. We designed a panel of five 40 bp random sequence dsRNAs with GC content varying from 0% to 100% and used them to test whether dsRADAR binding is influenced by base composition similar to J2 and K1 (Fig. 4, Supplementary Fig. S1b). For both 1x and 4x dsRADAR constructs, dsRNAs having intermediate GC content (25-75%) showed comparable affinities (150-200nM) (Fig. 4, Supplementary Table. S2). At 100% GC content, both 1x and 4x dsRADARs displayed a slightly reduced binding affinity (1x *K*d = 320 nM, 4x *K*d = 480 nM), while 4x dsRADAR exhibits a slightly higher apparent affinity with 0% GC substrate (*K*d = 62 nM).

**Figure 4.**
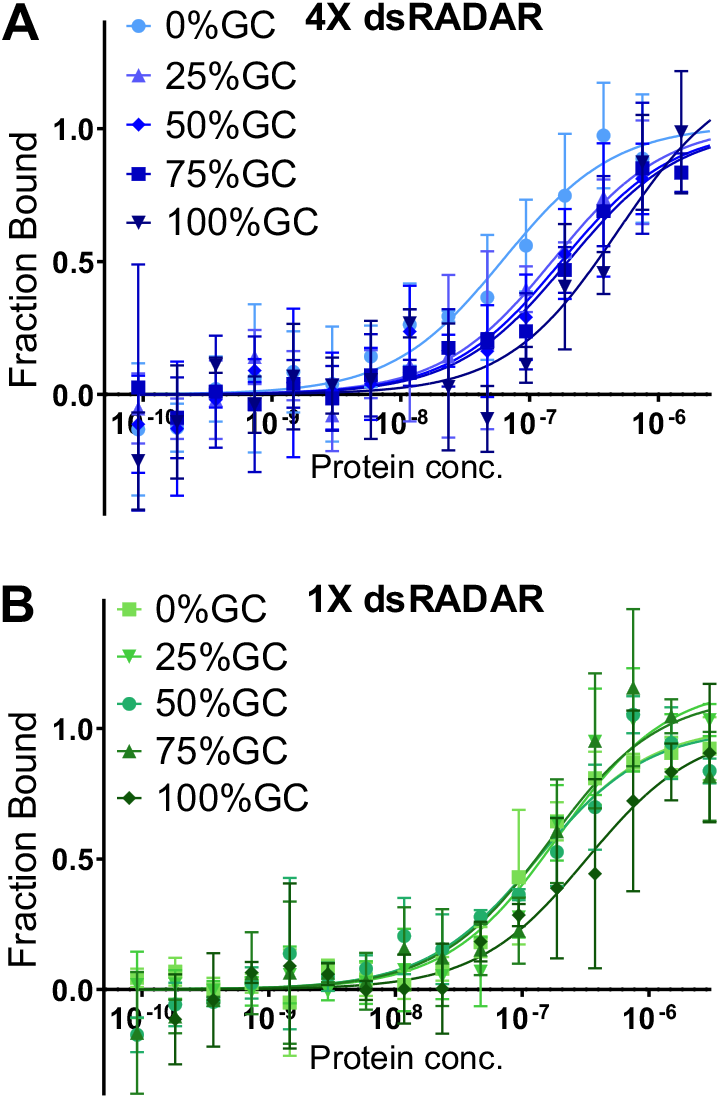
dsRADARs bind dsRNA in a nearly sequence agnostic manner. MST binding curves of 40 bp dsRNA having varying GC-content with (a) 4x dsRADAR and (b) 1x dsRADAR. All MST experiments were performed in 20 mM Tris, pH=7.6, 125 mM NaCl, 30 mM KCl, 5 mM MgCl_2_, 0.5 mM DTT.

This trend is consistent with the structural properties of RNA duplexes. Because GC-rich duplexes form tighter and more rigid helices due to the additional hydrogen bond between G-C base pairs and enhanced base stacking, the resulting decrease in helical flexibility may disrupt optimal duplex backbone interactions with dsRADARs and other dsRNA binding proteins.^29,40^ Conversely, AU-rich duplexes are thermodynamically weaker and more flexible, potentially facilitating the interaction with dsRADAR. Compared to the state-of-the-art J2 antibody, which completely loses binding affinity for dsRNAs having GC content above 63%, dsRADAR only exhibits a modest affinity variation from 0% to 100% GC content.^29^ Together, these results demonstrate that dsRADARs are able to tolerate variation in thermodynamic stability and helix rigidity so long as the dsRNA still maintains an overall A-form helical structure.

As a potential tool to detect, image, and enrich dsRNA of cellular and viral origins, it is important for dsRADARs to selectively bind dsRNA over other biologically relevant nucleic acids present in cells. We used MST to evaluate the interaction of both dsRADAR constructs with dsRNA, dsDNA, ssRNA, yeast tRNA, and RNA-DNA hybrid duplexes to identify any potential non-specific binding interactions that could complicate the interpretation of cell-based data (Fig. 5). Even though 4x dsRADAR can recognize short (≥8bp) dsRNA duplexes, both constructs still robustly discriminate against tRNA, indicating that dsRADARs do not promiscuously recognize compact and highly folded RNA. Both dsRADARs also showed high selectivity against dsDNA, ssRNA, and even RNA-DNA hybrid duplexes, which adopt an A/B intermediate conformation that closely resembles A-form dsRNA. These results demonstrate that dsRADARs only recognize a fully canonical A-form RNA duplex and have no measurable cross-reactivity to other common nucleic acid structures that may be encountered during biological assays.

**Figure 5.**
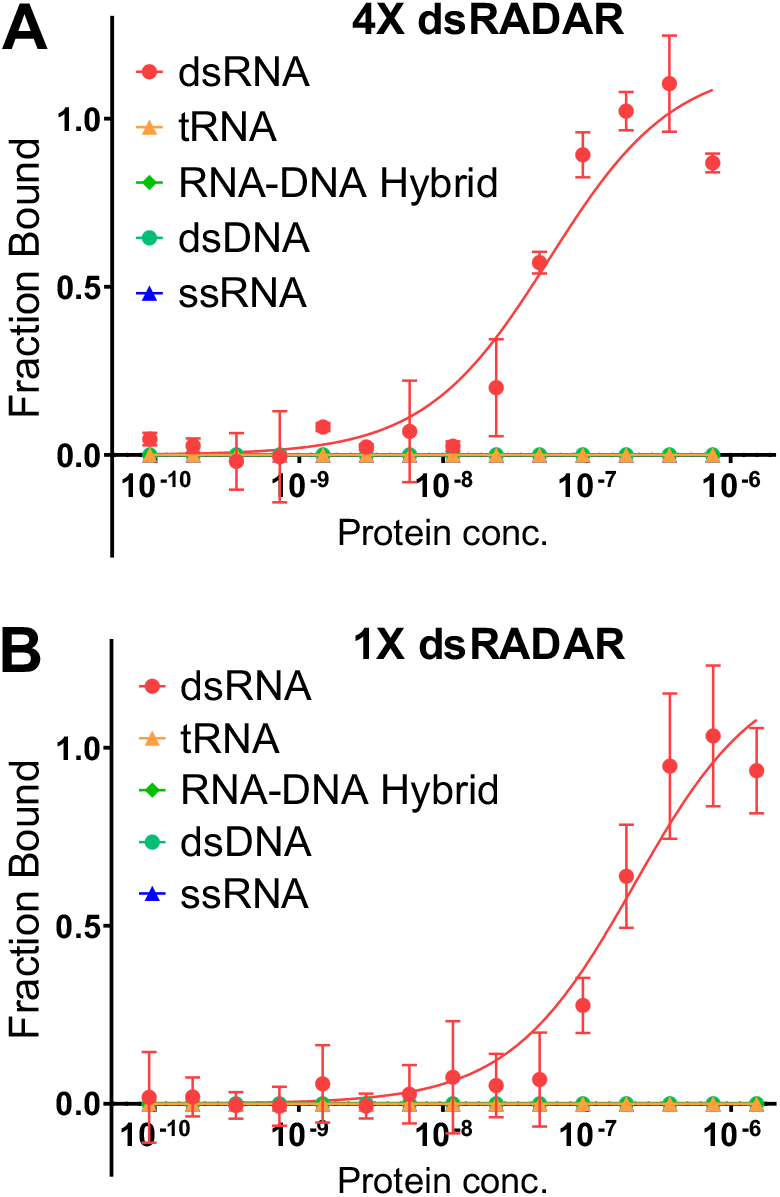
dsRADARs bind dsRNA with high specificity. MST binding curves of off-target substrates tRNA, RNA-DNA hybrid, dsDNA and ssRNA with (a) 4x dsRADAR and (b) 1x dsRADAR. All MST experiments were performed in 20 mM Tris, pH=7.6, 125 mM NaCl, 30 mM KCl, 5 mM MgCl_2_, 0.5 mM DTT.

### 4. dsRADARs recognize dsRNA accumulation in viral infected cell-based assays

Human astroviruses (HAstVs) are positive-sense, single-stranded RNA viruses that represent a major cause of pediatric gastroenteritis worldwide.^41,42^ HAstVs are classified into two main groups: classical (serotypes 1-8) and novel strains, with classical HAstVs accounting for up to 10% of sporadic cases of non-bacterial diarrhea in children.^41–43^ Previous studies have shown that positive-sense RNA viruses generate dsRNA as replication intermediates that can be detected by dsRNA-specific antibodies.^6^ To evaluate the ability of dsRADAR to detect dsRNA under physiological conditions, we infected Caco-2 cells with HAstV1 and performed immunofluorescence imaging (Fig. 6a-b). Compared to uninfected controls, HAstV1-infected cells exhibited an increased fluorescence signal detected by both dsRADARs and dsRNA antibodies (J2 and K1) (Fig. 6a-d). Furthermore, infected cells treated with RNase III to selectively digest dsRNA resulted in a substantial reduction in dsRADAR fluorescence signal, validating the specificity of dsRADARs for dsRNA (Fig. 6e). We note that 1x and 4x dsRADAR reveal punctate dsRNA clusters localized in the perinuclear region and which are colocalized with viral capsid protein. This observed localization is consistent with previous reports that HAstV replication occurs within double-membrane vesicles (DMVs) formed near the endoplasmic reticulum, which serve as sites for viral RNA replication.^44^ These observations indicate that the dsRADAR signal corresponds to the accumulation of viral dsRNA replication intermediates.

**Figure 6.**
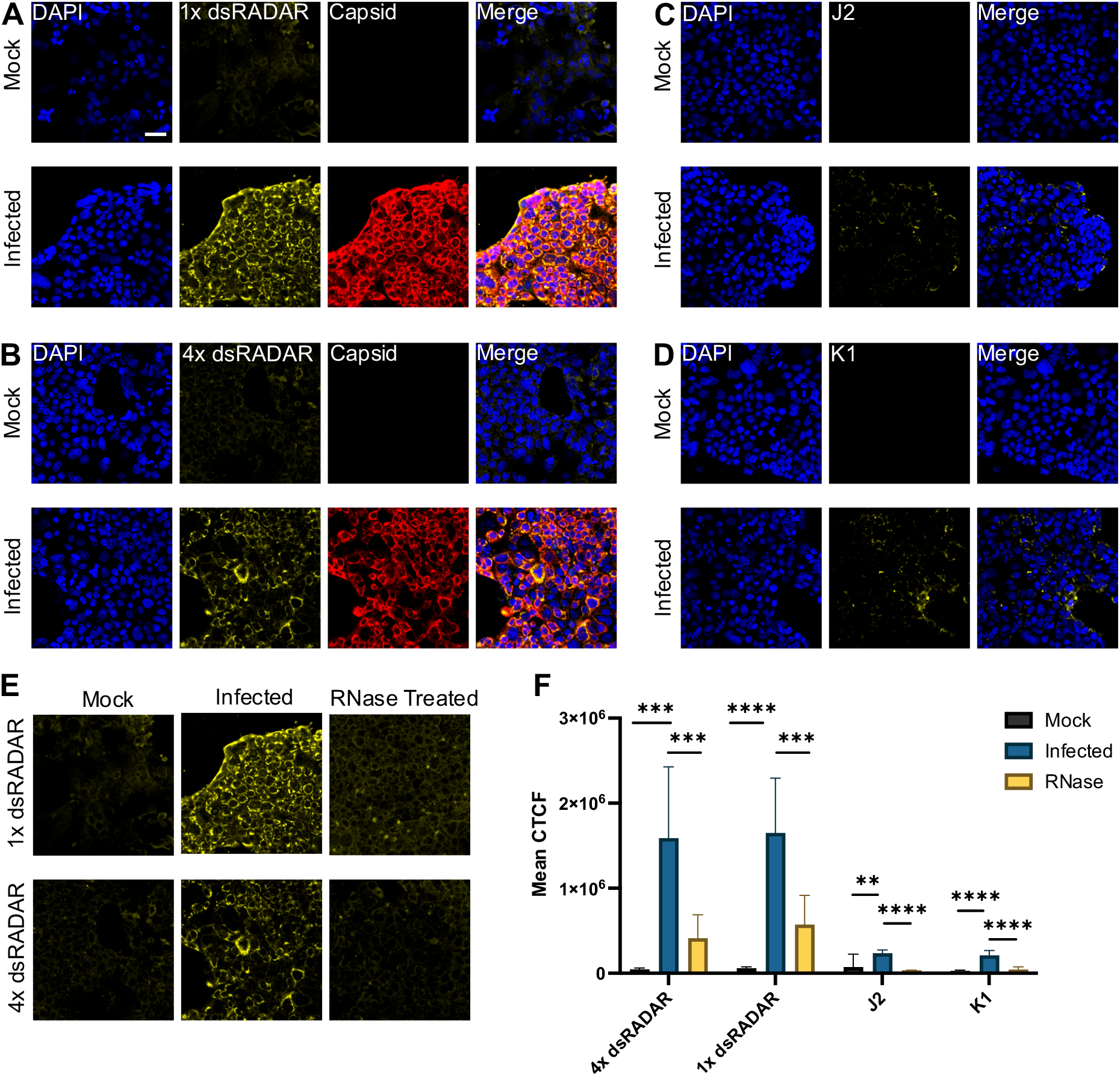
dsRADAR offers improved detection of viral dsRNA in cell-based infection assays. Compared with uninfected cells (*top*), Infected cells (*bottom*) show dsRNA accumulation with (a) 1x dsRADAR (*yellow*) and (b) 4x dsRADAR (*yellow*) colocalizing with HAstV1 specific capsid (*red*). Representative immunofluorescence images of uninfected cells (*top*) and infected cells with (c) J2 dsRNA antibody (*yellow*) and (d) K1 dsRNA antibody (*yellow*). (e) Immunofluorescence images after RNase III treatment of infected cells (*right*) compared with untreated infected cells (*middle*) and mock cells (*left*). (f) Quantification of immunofluorescence of mock cells, HAstV1-infected cells, and RNase treated HAstV1-infected cells. Statistical significance was determined using repeated-measures one-way ANOVA with Geisser-Greenhouse correction followed by Šídák’s multiple comparisons test. **P < .0021; ***P < .0002; ****P < .0001; n.s., not significant (P > .05). Scale bar, 50 *μ*m.

Quantitative analysis of fluorescence intensities demonstrates that both dsRADARs produce a stronger fluorescence signal in infected cells compared to dsRNA antibodies, with no appreciable increase in background fluorescence, indicating a drastically increased signal-to-noise for the detection of dsRNA accumulation. This enhanced signal was also achieved using a substantially lower affinity reagent concentration of 5 nM for dsRADAR compared to 33 nM for J2 and K1 antibodies. This improved performance reflects both the broader detection range and higher sensitivity of dsRADARs relative to conventional dsRNA antibodies. Furthermore, single-cell linear regression analysis between viral capsid signal and dsRADAR fluorescence (Supplementary Fig. S6) revealed a strong positive correlation, indicating that dsRADAR signal increases with HAstV1 level. Together, these results reveal that the dsRADAR constructs not only sensitively detect viral dsRNA in the cellular context but also quantitatively report on the extent of viral infection.

### 5. dsRADARs can detect endogenous dsRNA in FFPE tissue samples

Under physiological conditions, healthy human cells contain low levels of endogenous dsRNA that arise from sources such as structured noncoding RNAs, RNA interference pathways, and transcripts from repetitive genomic elements.^4^ However, in pathological contexts, dysregulation of RNA processing can lead to the accumulation of endogenous dsRNA, even in the absence of viral infection.^18–20^ An important regulator of cellular dsRNA metabolism is ADAR1, an RNA deaminase that catalyzes adenosine-to-inosine editing within endogenous long dsRNA regions, preventing the activation of innate cellular immune responses by structurally disrupting the duplex.^45^ Halstead et al. have recently demonstrated that the specific deletion of ADAR1 and MAVS from gastric parietal cells (Adar^ΔPC^ ; Mavs^-/-^) leads to gastric epithelial dsRNA accumulation, as monitored by K1 dsRNA antibody staining.^46^

To assess whether dsRADAR can detect pathologically-relevant accumulation of endogenous dsRNA, we performed immunofluorescence staining on formalin-fixed paraffin-embedded (FFPE) gastric tissue sections from Adar^ΔPC^, Mavs^-/-^ mouse models. As a control, we included the Tlr3^-/-^, Adar1^F/+^ mouse model, in which ADAR1 is intact in parietal cells and retains the same expression level as wild type (Fig. 7). Both the 1x and 4x dsRADAR constructs are able to detect substantial cytosolic dsRNA accumulation in the Adar^ΔPC^, Mavs^-/-^ tissue, whereas control samples showed minimal signal. We note that the dsRNA signal appears as discrete cytosolic puncta rather than being diffusely distributed. This suggests that the clustered localized dsRNA could be derived from compartmentalized sources of dsRNA, such as mitochondria. This observation aligns with previous studies utilizing dsRNA immunoprecipitation sequencing (dsRIP-seq) analysis indicating that the dsRNA accumulated in Adar^ΔPC^, Mavs^-/-^ are enriched for mitochondrial dsRNA.^46^ Together, these data validate that dsRADARs are compatible with FFPE tissue histochemistry and sensitive to the accumulation of endogenous dsRNA in disease-relevant model tissues.

**Figure 7.**
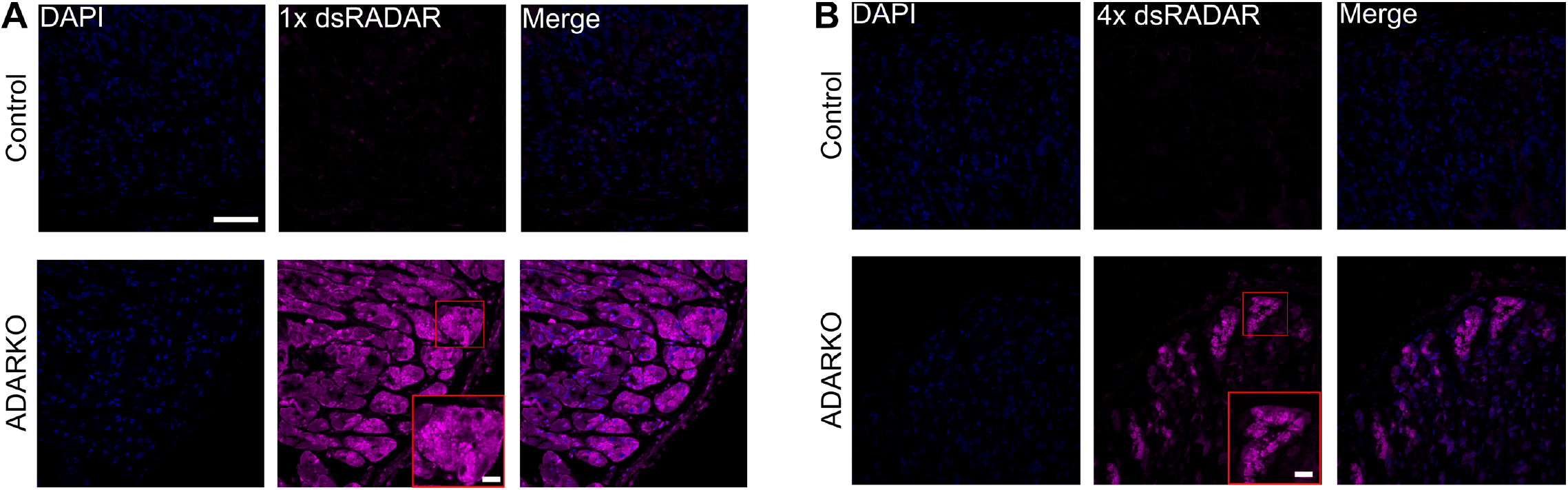
dsRADAR can detect endogenous dsRNA in FFPE tissue samples. Representative confocal images of Adar^ΔPC^; Mavs^-/-^ tissue (*bottom; ADARKO*) immunostained with (a) 1x dsRADAR (*magenta*) and (b) 4x dsRADAR (*magenta*). Tlr3^-/-^; Adar1^F/+^ mouse model was used as control (*top*). Scale bar, 50 *μ*m. Scale bar of zoomed in section, 10 *μ*m.

## Discussion

Double-stranded RNA is a critical signaling molecule to alert cells to viral infection, but can also cause harmful effects when dysregulated in disease or present as a contaminant from production of RNA vaccines and therapeutics. Despite this importance, tools to visualize and quantify dsRNA remain limited. Developing robust affinity reagents for dsRNA is particularly challenging because it is not a discrete molecular entity as is the case with small molecules or proteins. Rather, dsRNA can have essentially infinite variation in its specific nucleotide sequence and can vary widely in the length and finer secondary structure of the duplex region. The most widely used reagents, monoclonal antibodies J2 and K1, were developed to detect dsRNA in cells.^27^ However, J2 is incapable of binding to sequences having high GC content or short duplex regions and K1 has a strong preference for poly I:C, which is not representative of cellular dsRNA. These sequence and structure biases undermine the reliability of the available antibodies, fundamentally constraining their utility for rigorous quantification of dsRNA in cells.^27–29^ We hypothesized that nature could provide a superior solution to detecting dsRNA as many dsRBDs have been evolved for precisely the desired task – unbiased and sensitive detection of cellular dsRNA. In particular, we leveraged the molecular recognition capabilities of the dsRNA-binding domains of ADAR enzymes, as these enzymes are known to be involved in cellular regulation of dsRNA. Through systematic screening of ADAR dsRBDs and engineering of domain and linker architectures, we were able to develop dsRADARs as a set of improved affinity reagents that enable robust, sequence-independent, and highly sensitive recognition of cellular dsRNA.

To develop our dsRADAR reagents, we first systematically screened the dsRBDs of all three human ADARs to identify the most suitable candidate for unbiased dsRNA recognition. Although the RNA editing preferences of ADAR1 and ADAR2 are thought to be mainly determined by the deaminase domain, our results show that their isolated dsRBDs can have highly variable binding to randomized dsRNA substrates.^35–38^ This indicates that the substrate selectivity in ADAR1 and ADAR2 arises not only from the enzyme active sites but also from their dsRBD regions. In contrast, although ADAR3 is evolutionarily conserved and shares substantial sequence and structural homology with other members of the ADAR family, its dsRBDs displayed strong dsRNA affinity to both of the initially tested randomized dsRNA substrates, making it the optimal starting point for a dsRNA detection tool.^30^

By dissecting the contribution of each ADAR3 dsRBD and swapping domains, we show that robust binding requires cooperative interaction between dsRBD1 and dsRBD2. Neither tandem domain repeat construct nor any of the individual domains reproduced the affinity of the native tandem domains, demonstrating that both domains contribute distinctly but complementarily to the interactions that are necessary for high-affinity dsRNA binding. Linker engineering further revealed that interdomain spacing and flexibility are significant factors for dictating the binding mode with dsRNA, thus modulating the duplex length required for recognition. Specifically, we discovered that connecting the dsRBDs with a 4x (80 aa) flexible linker resulted in a minimal 8 bp dsRNA length for binding whereas replacement with a shorter 1x (20 aa) linker increased the minimum bound duplex size to 25 bp. The shorter 8 bp motif is consistent with the binding footprint of a single dsRBD, leading us to hypothesize that the longer linker enables the dsRBDs to bind cooperatively in a “clamp” mode on opposite faces of the same duplex region while the shorter linker is not able to accommodate this conformation and instead adopts an “in series” binding conformation in which each dsRBD binds to a separate dsRNA segment (Fig.3).

The sensitivity and flexibility of dsRADARs highlight several major improvements over existing antibody-based tools. For example, the antibodies have a fixed dsRNA recognition cutoff of 30-40 bp, which limits the detection of physiologically important short blunt-end duplexes that could serve as RIG-I ligands. In contrast, 4x dsRADAR readily binds these minimal dsRNA features in addition to the longer dsRNA duplexes that J2 also detects.^5,7,30,34–39,47,48^ Moreover, depending on the sensitivity required and types of dsRNA to be detected, the dsRADAR reagents can be used individually or in concert to specifically detect dsRNAs of different lengths. In the future, we envision elaborating these reagents into proximity-based sensors in which two dsRADAR reagents must bind adjacent to one another to give rise to a signal, as this would allow further tuning of the size cutoff up to 50 bp. In addition to greater flexibility in substrate length requirements, dsRADARs demonstrate nearly sequence agnostic binding. In particular, dsRADARs are able to robustly bind dsRNAs spanning 0-100% GC content, while J2 shows reduced affinity for dsRNA with increasing GC content and no detectable binding above 63% GC content.^29^ This is highly important for ensuring rigor and reproducibility, as 4x dsRADAR is able to recognize biologically relevant but previously difficult or impossible to detect substrates such as short viral replication intermediates of negative-strand viruses and GC-rich RNAs, including Alu repeats (averaging 62.7% GC) and transcribed CpG islands (by definition, regions of 200 bp or more having ≥50% GC content).^49,50^ The ability to detect these dsRNAs in addition to the more common mixed GC content, long dsRNAs provides a more accurate picture of the dsRNA landscape in a cell, leading to more reliable biological insights. Additionally, the high sensitivity and small binding footprint enables dsRADARs to bind densely along dsRNA, significantly enhancing signal-to-background ratio and doing so at lower concentrations than are required for the antibodies. The longer duplex requirement of 1x dsRADAR can also be used advantageously in selectively discriminating between short or transient duplex regions that may fall within the detection range of the 4x construct.

While the main goal of our *in vitro* characterization studies was to generate an improved affinity reagent for dsRNA, our discovery of the relatively sequence agnostic binding of dsRADARs also has important implications for the biological function of its native parent protein, ADAR3. The deaminase domain of human ADAR3 is known to be catalytically dead and thus the protein has been proposed to regulate posttranscriptional modification by acting as a competitor with ADAR1 and ADAR2 at specific sites to suppress A-to-I editing. ADAR3-specific targets have been found to partially overlap with subsets of ADAR1 and ADAR2 substrates, though it also has unique known targets distinct from reported ADAR1 and ADAR2 substrates.^39,48^ This demonstrates that full-length ADAR3 and its multimeric forms may have intrinsic sequence preferences and ability to selectively recognize particular RNAs. Our observation that the isolated dsRBDs of ADAR3 bind dsRNA with a high level of sequence tolerance suggests that any transcript selectivity observed *in vivo* is likely controlled predominantly through the noncatalytic deaminase domain and/or protein-protein interactions of ADAR3.

While our *in vitro* characterization data proved encouraging and insightful, we were most excited to observe the performance of dsRADARs for immunostaining in disease-relevant cell and tissue samples. In cell-based assays, dsRADARs were able to detect virus-derived dsRNA in HAstV1-infected Caco-2 cells, producing robust signals that correlated with infection status and viral protein expression with greater signal-to-noise than either the J2 or K1 antibodies. This performance extends further to complex tissue samples, as dsRADARs effectively recognized endogenous dsRNA accumulation in Adar^ΔPC^, Mavs^-/-^ FFPE mouse gastric tissue. The ability of both dsRADAR constructs to maintain binding activity in cell and preserved tissue samples validates their reliability and compatibility with standard histological workflows, supporting dsRADAR as a broadly applicable dsRNA probe.

Together, these findings establish dsRADARs as highly sensitive dsRNA detection tools that are biologically compatible with standard laboratory workflows and overcome the limitations of existing dsRNA antibodies. Their minimal sequence bias increases rigor and reproducibility in cellular experiments, and their high binding affinity and flexible detection range provide advantages for studying diseases involving accumulation of both viral and endogenous dsRNA. Looking to the future, dsRADARs can be easily adapted for ELISA based quantification of dsRNA contaminants in RNA vaccines and therapeutics. Additionally, unlike antibodies, the dsRADAR constructs can be genetically encoded and easily equipped with myriad tags, recruitment modules, or effector proteins, opening the door to a wide range of potential live cell technologies, and research in these areas is ongoing in our lab.

## Supporting Information

Experimental materials and methods including supplementary table and supporting figures (PDF)

## Acknowledgements

This work was supported by the National Institutes of Health (R35GM144075 to J.M.H.). The content is solely the responsibility of the authors and does not necessarily reflect the official views of the National Institute of Health. We wish to thank MilliporeSigma in St. Louis for insightful conversations and helpful discussions. We would like to thank Dr. Alexandria L. Quillin as well for her input in analysis of cell and tissue immunofluorescence staining. We would also like to thank Somya Aggarwal for helpful discussion.

