## Supplemental Information for "dsRADAR: Imaging and Quantifying Cellular dsRNA by Repurposing RNA Binding Proteins"

**Affiliations:**
[1]Department of Chemistry, Washington University in St. Louis, MO, USA.
[2]Division of Infectious Diseases, Department of Medicine, Edison Family Center for Genome Sciences and Systems Biology, Washington University School of Medicine, St. Louis, MO, USA.
[3]Division of Gastroenterology, Department of Medicine, Washington University in St. Louis School of Medicine, St. Louis, MO, USA.
[4]Department of Molecular Microbiology, Washington University School of Medicine, St. Louis, MO, USA.
[5]Department of Molecular Cell Biology, Washington University in St. Louis School of Medicine, St. Louis, MO, USA.

### Supplemental Information

#### Table of Contents

**Materials and Methods** S1-S5

**Table S1.** RNA substrate sequence S6

**Table S2.** Summary of $k_d$ values measured by MST S7

**Table S3.** Plasmid of dsRADAR constructs S8

**Figure S1.** RNA substrate structure S9

**Figure S2.** Validation of RNA substrate labeling by RNA gel electrophoresis S10

**Figure S3.** Characterization of dsRBD binding to random sequence RNAs: MST binding curves showing ΔFnorm responses S11

**Figure S4.** *In vitro* characterization of dsRADAR: MST binding curves showing ΔFnorm responses S12

**Figure S5.** Immunofluorescence analysis of infected cells after RNase treatment stained with J2 and K1 S13

**Figure S6.** Linear regression analysis of viral capsid signal and dsRADAR staining S14

**Figure S7.** Optimization of dsRADAR concentration for tissue staining S15

### Materials and Methods:

#### Safety Statement

No unexpected or unusually high safety hazards were encountered in this study.

#### Materials

Methanol, Triton X-100, Tween 20, tris hydrochloride, sodium chloride, sodium peroxide, imidazole, magnesium chloride, potassium chloride, and DTT were purchased from Sigma Aldrich. Nuclease-free water, ultra-pure glycerol, Alexa Fluor 555, Alexa Fluor 647, poly-D-Lysine, Hoechst 33342, and 96-well plates were purchased from Thermo Fisher Scientific.

#### RNA Preparation

All RNA-related procedures were performed using RNase-free conditions. Work surfaces and equipment were treated with RNaseZap (Ambion), and all buffers were prepared with nuclease-free water. Fluorescently labeled hairpin RNAs containing 10 bp or 20 bp duplex regions were obtained from the DNA/Peptide Core Facility at the University of Utah. Longer dsRNA substrates and RNA substrates of varying GC content were produced by in vitro transcription (IVT). DNA templates were ordered from IDT and transcribed using the HiScribe T7 High Yield RNA Synthesis Kit (NEB), followed by DNase treatment and purification using the Monarch RNA Cleanup Kit (NEB). RNA integrity and size were verified on 10% denaturing urea-PAGE gels. For 3'-end fluorescent labeling, purified RNAs were first oxidized with 10 mM sodium periodate at room temperature for 90 min, then incubated with Cy5 hydrazide (Lumiprobe) for 2.5 h at 37 ˚C. Excess dye was removed by ethanol precipitation, and labeling efficiency was assessed spectroscopically using a NanoDrop spectrophotometer. Yeast tRNA was purchased from Thermo Fisher Scientific.

#### Protein Expression and Purification

Recombinant dsRBD constructs (6xHis-MBP-TEV-dsRBD) were cloned into a pET vector and transformed into E. coli BL21(DE3) cells. Overnight seed cultures were grown at 37 °C and used to inoculate 500 mL LB medium. Cultures were grown at 37 °C for ~4 h, and protein expression was induced with 0.5 mM IPTG. Induced cultures were incubated at 18 °C for 16-18 h and harvested by centrifugation (5,000 x g, 10 min, 4 °C). Cell pellets were resuspended in lysis buffer (GoldBio) according to the manufacturer's instructions. Lysates were clarified by centrifugation at 16,000 x g for 25 min at 4 °C, and the supernatant was incubated with 2 mL cobalt affinity resin (GoldBio) pre-equilibrated in Buffer A (10 mM Tris-HCl pH 7.5, 200 mM NaCl, 10 mM imidazole). Binding was performed by gentle rotation at 4 °C for 1.5 h. The resin was washed twice by resuspension in 50 mL Buffer A followed by centrifugation (3,000 x g, 2 min, 4 °C). Washed resin was packed into a gravity column and washed sequentially with 3 column volumes of Buffer A containing 50 mM and then 100 mM imidazole. Bound proteins were eluted stepwise with Buffer A supplemented with 150 mM, 200 mM, and 300 mM imidazole. Eluted fractions were analyzed by SDS-PAGE, pooled and buffer-exchanged into storage buffer (20 mM Tris-HCl pH 7.5, 200 mM NaCl, 5% glycerol, 1 mM DTT). Purified protein was flash-frozen in liquid nitrogen and stored at -80 °C.

**Microscale Thermophoresis**

MST measurements were performed on a Monolith NT.115 instrument (NanoTemper Technologies). RNA stocks were prepared in Binding Buffer (20 mM Tris-HCl pH 7.0, 50 mM NaCl, 60 mM KCl, 10 mM $MgCl_2$). Protein stocks were prepared in 20 mM Tris-HCl pH 8.0, 200 mM NaCl, 1 mM DTT. RNA substrates were annealed by heating to 75 °C for 2 min and slowly cooling to 4 °C over 30 min. Serial dilutions of protein were mixed with a constant concentration of Cy5-labeled RNA at a 1:1 ratio, incubated 30 minutes on ice, and loaded into premium capillaries. Measurements were conducted in triplicate following the manufacturer's recommended settings.

**Viral Infection and Immunostaining**

Cells were cultured at 37 °C in a humidified incubator with 5% CO2. For immunofluorescence experiments, black 96-well plates (Cellvis) were coated in poly-D-lysine (Gibco) following the manufacturer's protocol. 2.5 x $10^4$ Caco2 cells were seeded in 96-well plates at 72 h before infection. Cells were infected with HAstV1 at MOI = 10 for 1 h at 37 °C, followed by 2 washes with PBS. Fresh media were added, and the cells were incubated at 37 °C until fixation. Immunostaining workflow was performed based on general protocol outlined by Quillin et.al.[1] Cells were then fixed in 100% methanol and incubated at 4 °C for 15 minutes. Wells were washed twice with 1X TBS. Cells were then incubated with a 0.1% Triton X-100 in 1X TBS permeabilization solution at room temperature for 10 minutes followed by two washes in 1X TBS. Cells were then incubated in a blocking solution containing 3% bovine serum albumin, 0.1% Triton X-100, and 0.1% Tween 20 in 1X PBS for 1 hour at room temperature. Wells were then washed twice with 1X PBS. Cells were incubated with 5 nM dsRADAR containing blocking buffer at room temperature for 1 hour. Cells were then washed three times with 1X TBS, incubating for 5 minutes each time. Cells were then incubated for 1 hour at room temperature with a 1:1000 anti-MBP antibody (New England BioLabs) solution diluted in blocking buffer. Cells were washed three times, with 5-minute incubation periods. Finally, a staining solution containing 1:1000 goat anti-mouse Alexa Flour 647 and 1:1000 Hoechst nuclear dye in calcium containing blocking buffer was incubated in the wells for 1 hour at room temperature in the dark. Cells were then washed three times for 5 minutes each time. Cells were then imaged. All immunostaining incubations were completed with 200 mL volumes and solutions were prepared fresh for each experiment.

**FFPE Tissues Immunofluorescence**

Immunostaining workflow inspired by Quillin et.al.[2] Slides were first deparaffinized and then rehydrated through a graded ethanol series followed water. Slides were then washed in 1X TBS for 5 minutes at room temperature with gentle rocking. Antigen retrieval was performed using sodium citrate buffer in a pressure cooker. Slides were then washed in 1X TBS for 15 minutes at room temperature. Tissues were stained as previously described for cells with 2nM dsRADAR. Following staining, excess buffer was removed, and a small drop of antifade mounting medium was applied to a coverslip and placed on the tissue section. Slides were allowed to cure overnight at room temperature in the dark and imaged the following day. Mounted slides were stored at 4 °C in a dark slide box. All incubations were performed under nuclease-free conditions in a humidified chamber. Solutions were prepared fresh for each experiment.

**Microscopy and Image Analysis**

Stained cells and tissues were imaged using a Nikon Spinning Disk for widefield microscopy with a 20x air objective and confocal microscopy with a 60x oil objective. Laser excitation at 405 nm was used to image Hoechst 33342; excitation at 640 nm was used to image Alexa Fluor 647; excitation at 560 nm was used to image Alexa Fluor 555. Gain and exposure settings for each laser were optimized to achieve sufficient fluorescence and minimize oversaturation. The resulting images were analyzed to determine the fluorescence of each cell using FIJI. Fluorescence was quantified using mean fluorescence or the Corrected Total Cellular Fluorescence (CTCF). The CTCF was calculated by measuring the area and integrated density and using the following equation: CTCF=integrated density−(cell area × mean background fluorescence) CTCF=integrated density−(cell area × mean background fluorescence).

**Table S1. RNA substrate sequences.**

| RNA Substrate | Sequence (3' to 5') |
|---|---|
| rsRNA-1 (20bp) | GUGCAGUGGCGCAAUCUUGGUUGUUCCAAGAUUGCGCCACUGCAC |
| rsRNA-2 (25bp) | GGGAUCAAUAUGCUAAGCGAUCCCUUGUUGGGAUCGCUUAGCAUAUUGAUCCC |
| 6bp Hairpin | GGAUUGUUCGCAAUCC |
| 8bp Hairpin | GGAUCAUGUUCGCAUGAUCC |
| 10bp Hairpin | GGAUCAUCUGUUCGCAGAUGAUCC |
| 20bp Hairpin | GUGCAGUGGCGCAAUCUUGGUUGUUCCAAGAUUGCGCCACUGCAC |
| 25bp Hairpin | GGGAUCAAUAUGCUAAGCGAUCCCUUGUUGGGAUCGCUUAGCAUAUUGAUCCC |
| 30bp ssRNA (+) | GGGAUAAUUUGUAAAGAAGGUCAUUAUCCC |
| 30bp ssRNA (-) | GGGAUAAUGACCUUCUUUACAAAUUAUCCC |
| 40bp ssRNA (+) | GGGAUAAUUUGUAAAGAAGGUCAAUAUUGUACAUUAUCCC |
| 40bp ssRNA (-) | GGGAUAAUGUACAAUAUUGACCUUCUUUACAAAUUAUCCC |
| 50bp ssRNA (+) | GGGAUAAUUUGUAAAGAAGGUCAAUAUUGUACCAAACUUGAUAUUAUCCC |
| 50bp ssRNA (-) | GGGAUAAUAUCAAGUUUGGUACAAUAUUGACCUUCUUUACAAAUUAUCCC |
| 0% GC ssRNA (+) | GUAAUAUUAUUAUAAUUUUAUUUUAUUAAAAUAUUAAUUC |
| 0% GC ssRNA (-) | GAAUUAAUAUUUUAAUAAAAUAAAAUUAUAAUAAUAUUAC |
| 25% GC ssRNA (+) | GUGACAUUGUUACAAUGUUAGUUUGUUAGAAUGUUGAUUC |
| 25% GC ssRNA (-) | GAAUCAACAUUCUAACAAACUAACAUUGUAACAAUGUCAC |
| 50% GC ssRNA (+) | GUGCCAGUCUCAGACUGUGACUCUGUGAGACUGCUGACUC |
| 50% GC ssRNA (-) | GAGUCAGCAGUCUCACAGAGUCACAGUCUGAGACUGGCAC |
| 75% GC ssRNA (+) | GAGUCAGGACCCUCGCACCGCCACCGUCUGAGGCUGGGAC |
| 75% GC ssRNA (-) | GUCCCAGCCUCAGACGGUGGCGGUGCGAGGGUCCUGACUC |
| 100% GC ssRNA (+) | GGGCCCGGGCCCGCGCGCCGGCGCCGGGCCCGGCCGGGCC |
| 100% GC ssRNA (-) | GGCCCGGCCGGGCCCGGCGCCGGCGCGCGGGCCCGGGCCC |

**Table S2. Summary of *kd* values measured by MST.**

| RNA Substrate | $K_d$ value of 4x dsRADAR | $K_d$ value of 1x dsRADAR |
|---|---|---|

| | | |
|---|---|---|
| 6bp Hairpin | N/A | N/A |
| 8bp Hairpin | 110nM ± 70nM | N/A |
| 10bp Hairpin | 130nM ± 70nM | N/A |
| 20bp Hairpin | 54nM ± 20nM | N/A |
| 25bp Hairpin | 67nM ± 30nM | 130nM ± 200nM |
| 30bp dsRNA | 260nM ± 230nM | 320nM ± 140nM |
| 40bp dsRNA | 190nM ± 150nM | 180nM ± 140nM |
| 50bp dsRNA | 220nM ± 100nM | N/A |
| 0% GC dsRNA | 62nM ± 64nM | 130nM ± 50nM |
| 25% GC dsRNA | 150nM ± 190nM | 190nM ± 80nM |
| 50% GC dsRNA | 180nM ± 290nM | 130nM ± 76nM |
| 75% GC dsRNA | 200nM ± 200nM | 160nM ± 100nM |
| 100% GC dsRNA | 480nM ± 470nM | 320nM ± 400nM |
| | **$K_d$ value of ADAR3 dsRBDs** | **$K_d$ value of ADAR3 Tandem dsRBD1** |
| rsRNA-1 | 170nM ± 80nM | 150nM ± 100nM |
| rsRNA-2 | 150nM ± 120nM | N/A |

**Table S3. Sequences of plasmids encoding ADAR variant constructs.** All sequences are shown in 3’ to 5’ direction. Sequence is annotated as following: Start Condon; 6xHN Tag; MBP Tag; TEV site; ADAR constructs.

**ADAR1 dsRBDs**

MGKYYHNHNHNHNHNHNSSGLVPRGSHMKIEEGKLVIWINGDKGYNGLAEVGKKFEKDTGIKVTVEHPDKLEEKFPQVAATGDG
PDIIFWAHDRFGGYAQSGLLAEITPDKAFQDKLYPFTWDAVRYNGKLIAYPIAVEALSLIYNKDLLPNPPKTWEEIPALDKELKAKGK
SALMFNLQEPYFTWPLIAADGGYAFKYENGKYDIKDVGVDNAGAKAGLTFLVDLIKNKHMNADTDYSIAEAAFNKGETAMTINGPW
AWSNIDTSKVNYGVTVLPTFKGQPSKPFVGVLSAGINAASPNKELAKEFLENYLLTDEGLEAVNKDKPLGAVALKSYEEELAKDPRI
AATMENAQKGEIMPNIPQMSAFWYAVRTAVINAASGRQTVDEALKDAQTSGGSSGSENLYFQGSGSSGGSMGSSHHHHHHSS
GLVPRGSHMKIEEGKLVIWINGDKGYNGLAEVGKKFEKDTGIKVTVEHPDKLEEKFPQVAATGDGPDIIFWAHDRFGGYAQSGLLA
EITPDKAFQDKLYPFTWDAVRYNGKLIAYPIAVEALSLIYNKDLLPNPPKTWEEIPALDKELKAKGKSALMFNLQEPYFTWPLIAADG
GYAFKYENGKYDIKDVGVDNAGAKAGLTFLVDLIKNKHMNADTDYSIAEAAFNKGETAMTINGPWAWSNIDTSKVNYGVTVLPTFK
GQPSKPFVGVLSAGINAASPNKELAKEFLENYLLTDEGLEAVNKDKPLGAVALKSYEEELAKDPRIAATMENAQKGEIMPNIPQMS
AFWYAVRTAVINAASGRQTVDEALKDAQTSGGSSGSENLYFQGSGSSGGSNPISGLLEYAQFASQTCEFNMIEQSGPPHEPRFK
FQVVINGREFPPAEAGSKKVAKQDAAMKAMTILLEEAKAKDSGKSEESSHYSTEKESEKTAESQTPTPSATSFFSGKSPVTTLLEC
MHKLGNSCEFRLLSKEGPAHEPKFQYCVAVGAQTFPSVSAPSKKVAKQMAAEEAMKALHGEATNSMASDNQPEGMISESLDNL
ESMMPNKVRKIGELVRYLNTNPVGGLLEYARSHGFAAEFKLVDQSGPPHEPKFVYQAKVGGRWFPAVCAHSKKQGKQEAADAA
LRVLIG*

**ADAR2 dsRBDs**

MGKYYHNHNHNHNHNHNSSGLVPRGSHMKIEEGKLVIWINGDKGYNGLAEVGKKFEKDTGIKVTVEHPDKLEEKFPQVAATGDG
PDIIFWAHDRFGGYAQSGLLAEITPDKAFQDKLYPFTWDAVRYNGKLIAYPIAVEALSLIYNKDLLPNPPKTWEEIPALDKELKAKGK
SALMFNLQEPYFTWPLIAADGGYAFKYENGKYDIKDVGVDNAGAKAGLTFLVDLIKNKHMNADTDYSIAEAAFNKGETAMTINGPW
AWSNIDTSKVNYGVTVLPTFKGQPSKPFVGVLSAGINAASPNKELAKEFLENYLLTDEGLEAVNKDKPLGAVALKSYEEELAKDPRI
AATMENAQKGEIMPNIPQMSAFWYAVRTAVINAASGRQTVDEALKDAQTSGGSSGSENLYFQGSGSSGGSQGAMGLPKNALMQ
LNEIKPGLQYTLLSQTGPVHAPLFVMSVEVNGQVFEGSGPTKKKAKLHAAEKALRSFVQFPNASEAHLAMGRTLSVNTDFTSDQA
DFPDTLFNGFETPDKAEPPFYVGSNGDDSFSSSGDLSLSASPVPASLAQPPLPVLPPFPPPSGKNPVMILNELRPGLKYDFLSESG
ESHAKSFVMSVVVDGQFFEGSGRNKKLAKARAAQSALAAIFNLHLDADPNSSS*

**ADAR3 dsRBDs**

MGKYYHNHNHNHNHNHNSSGLVPRGSHMKIEEGKLVIWINGDKGYNGLAEVGKKFEKDTGIKVTVEHPDKLEEKFPQVAATGDG
PDIIFWAHDRFGGYAQSGLLAEITPDKAFQDKLYPFTWDAVRYNGKLIAYPIAVEALSLIYNKDLLPNPPKTWEEIPALDKELKAKGK
SALMFNLQEPYFTWPLIAADGGYAFKYENGKYDIKDVGVDNAGAKAGLTFLVDLIKNKHMNADTDYSIAEAAFNKGETAMTINGPW
AWSNIDTSKVNYGVTVLPTFKGQPSKPFVGVLSAGINAASPNKELAKEFLENYLLTDEGLEAVNKDKPLGAVALKSYEEELAKDPRI
AATMENAQKGEIMPNIPQMSAFWYAVRTAVINAASGRQTVDEALKDAQTSGGSSGSENLYFQGSGSSGGSQGAMGAPKNALVQ
LHELRPGLQYRTVSQTGPVHAPVFAVAVEVNGLTFEGTGPTKKKAKMRAAELALRSFVQFPNACQAHLAMGGGPGPGTDFTSD
QADFPDTLFQEFEPPAPRPGLAGGRPGDAALLSAAYGRRRLLCRALDLVGPTPATPAAPGERNPVVLLNRLRAGLRYVCLAEPAE
RRARSFVMAVSVDGRTFEGSGRSKKLARGQAAQAALQELFDIQMPGADPNSSS*

**ADAR3 dsRBD1**

MGKYYHNHNHNHNHNHNSSGLVPRGSHMKIEEGKLVIWINGDKGYNGLAEVGKKFEKDTGIKVTVEHPDKLEEKFPQVAATGDG
PDIIFWAHDRFGGYAQSGLLAEITPDKAFQDKLYPFTWDAVRYNGKLIAYPIAVEALSLIYNKDLLPNPPKTWEEIPALDKELKAKGK
SALMFNLQEPYFTWPLIAADGGYAFKYENGKYDIKDVGVDNAGAKAGLTFLVDLIKNKHMNADTDYSIAEAAFNKGETAMTINGPW
AWSNIDTSKVNYGVTVLPTFKGQPSKPFVGVLSAGINAASPNKELAKEFLENYLLTDEGLEAVNKDKPLGAVALKSYEEELAKDPRI
AATMENAQKGEIMPNIPQMSAFWYAVRTAVINAASGRQTVDEALKDAQTSGGSSGSENLYFQGSGSSGGSQGAMGAPKNALVQ
LHELRPGLQYRTVSQTGPVHAPVFAVAVEVNGLTFEGTGPTKKKAKMRAAELALRSFVQ*

**ADAR3 dsRBD2**

MGKYYHNHNHNHNHNHNSSGLVPRGSHMKIEEGKLVIWINGDKGYNGLAEVGKKFEKDTGIKVTVEHPDKLEEKFPQVAATGDG
PDIIFWAHDRFGGYAQSGLLAEITPDKAFQDKLYPFTWDAVRYNGKLIAYPIAVEALSLIYNKDLLPNPPKTWEEIPALDKELKAKGK
SALMFNLQEPYFTWPLIAADGGYAFKYENGKYDIKDVGVDNAGAKAGLTFLVDLIKNKHMNADTDYSIAEAAFNKGETAMTINGPW
AWSNIDTSKVNYGVTVLPTFKGQPSKPFVGVLSAGINAASPNKELAKEFLENYLLTDEGLEAVNKDKPLGAVALKSYEEELAKDPRI
AATMENAQKGEIMPNIPQMSAFWYAVRTAVINAASGRQTVDEALKDAQTSGGSSGSENLYFQGSGSSGGSQGAMGPGERNPV
VLLNRLRAGLRYVCLAEPAERRARSFVMAVSVDGRTFEGSGRSKKLARGQAAQAALQELFDIQMPGADPNSSS*

**Tandem ADAR3 dsRBD1**

MGKYYHNHNHNHNHNHNSSGLVPRGSHMKIEEGKLVIWINGDKGYNGLAEVGKKFEKDTGIKVTVEHPDKLEEKFPQVAATGDG
PDIIFWAHDRFGGYAQSGLLAEITPDKAFQDKLYPFTWDAVRYNGKLIAYPIAVEALSLIYNKDLLPNPPKTWEEIPALDKELKAKGK

| |
|---|
| SALMFNLQEPYFTWPLIAADGGYAFKYENGKYDIKDVGVDNAGAKAGLTFLVDLIKNKHMNADTDYSIAEAAFNKGETAMTINGPW<br>AWSNIDTSKVNYGVTVLPTFKGQPSKPFVGVLSAGINAASPNKELAKEFLENYLLTDEGLEAVNKDKPLGAVALKSYEEELAKDPRI<br>AATMENAQKGEIMPNIPQMSAFWYAVRTAVINAASGRQTVDEALKDAQTSGGSSGSENLYFQGSGSSGGSQGAMGAPKNALVQ<br>LHELRPGLQYRTVSQTGPVHAPVFAVAVEVNGLTFEGTGPTKKKAKMRAAELALRSFVQGGGGSSGSETPGTSESATPESGGG<br>SSGSETPGTSESATPESGGGSSGSETPGTSESATPESGGGSSGSETPGTSESATPESAPKNALVQLHELRPGLQYRTVSQTGPV<br>HAPVFAVAVEVNGLTFEGTGPTKKKAKMRAAELALRSFVQA* |
| **Tandem ADAR3 dsRBD2** |
| MGKYYHNHNHNHNHNHNSSGLVPRGSHMKIEEGKLVIWINGDKGYNGLAEVGKKFEKDTGIKVTVEHPDKLEEKFPQVAATGDG<br>PDIIFWAHDRFGGYAQSGLLAEITPDKAFQDKLYPFTWDAVRYNGKLIAYPIAVEALSLIYNKDLLPNPPKTWEEIPALDKELKAKGK<br>SALMFNLQEPYFTWPLIAADGGYAFKYENGKYDIKDVGVDNAGAKAGLTFLVDLIKNKHMNADTDYSIAEAAFNKGETAMTINGPW<br>AWSNIDTSKVNYGVTVLPTFKGQPSKPFVGVLSAGINAASPNKELAKEFLENYLLTDEGLEAVNKDKPLGAVALKSYEEELAKDPRI<br>AATMENAQKGEIMPNIPQMSAFWYAVRTAVINAASGRQTVDEALKDAQTSGGSSGSENLYFQGSGSSGGSQGAMGPGERNPV<br>VLLNRLRAGLRYVCLAEPAERRARSFVMAVSVDGRTFEGSGRSKKLARGQAAQAALQELFDGGGGSSGSETPGTSESATPESG<br>GGSSGSETPGTSESATPESGGGSSGSETPGTSESATPESGGGSSGSETPGTSESATPESPGERNPVVLLNRLRAGLRYVCLAE<br>PAERRARSFVMAVSVDGRTFEGSGRSKKLARGQAAQAALQELFDIQMPGADPNSSS* |
| **dsRADAR construct with 1x synthetic linker** |
| MGKYYHNHNHNHNHNHNSSGLVPRGSHMKIEEGKLVIWINGDKGYNGLAEVGKKFEKDTGIKVTVEHPDKLEEKFPQVAATGDG<br>PDIIFWAHDRFGGYAQSGLLAEITPDKAFQDKLYPFTWDAVRYNGKLIAYPIAVEALSLIYNKDLLPNPPKTWEEIPALDKELKAKGK<br>SALMFNLQEPYFTWPLIAADGGYAFKYENGKYDIKDVGVDNAGAKAGLTFLVDLIKNKHMNADTDYSIAEAAFNKGETAMTINGPW<br>AWSNIDTSKVNYGVTVLPTFKGQPSKPFVGVLSAGINAASPNKELAKEFLENYLLTDEGLEAVNKDKPLGAVALKSYEEELAKDPRI<br>AATMENAQKGEIMPNIPQMSAFWYAVRTAVINAASGRQTVDEALKDAQTSGGSSGSENLYFQGSGSSGGSQGAMGAPKNALVQ<br>LHELRPGLQYRTVSQTGPVHAPVFAVAVEVNGLTFEGTGPTKKKAKMRAAELALRSFVQGGGGSSGSETPGTSESATPESPGER<br>NPVVLLNRLRAGLRYVCLAEPAERRARSFVMAVSVDGRTFEGSGRSKKLARGQAAQAALQELFDIQMPGADPNSSS* |
| **dsRADAR construct with 4x synthetic linker** |
| MGKYYHNHNHNHNHNHNSSGLVPRGSHMKIEEGKLVIWINGDKGYNGLAEVGKKFEKDTGIKVTVEHPDKLEEKFPQVAATGDG<br>PDIIFWAHDRFGGYAQSGLLAEITPDKAFQDKLYPFTWDAVRYNGKLIAYPIAVEALSLIYNKDLLPNPPKTWEEIPALDKELKAKGK<br>SALMFNLQEPYFTWPLIAADGGYAFKYENGKYDIKDVGVDNAGAKAGLTFLVDLIKNKHMNADTDYSIAEAAFNKGETAMTINGPW<br>AWSNIDTSKVNYGVTVLPTFKGQPSKPFVGVLSAGINAASPNKELAKEFLENYLLTDEGLEAVNKDKPLGAVALKSYEEELAKDPRI<br>AATMENAQKGEIMPNIPQMSAFWYAVRTAVINAASGRQTVDEALKDAQTSGGSSGSENLYFQGSGSSGGSQGAMGPGERNPV<br>VLLNRLRAGLRYVCLAEPAERRARSFVMAVSVDGRTFEGSGRSKKLARGQAAQAALQELFDGGGGSSGSETPGTSESATPESG<br>GGSSGSETPGTSESATPESGGGSSGSETPGTSESATPESGGGSSGSETPGTSESATPESPGERNPVVLLNRLRAGLRYVCLAE<br>PAERRARSFVMAVSVDGRTFEGSGRSKKLARGQAAQAALQELFDIQMPGADPNSSS* |

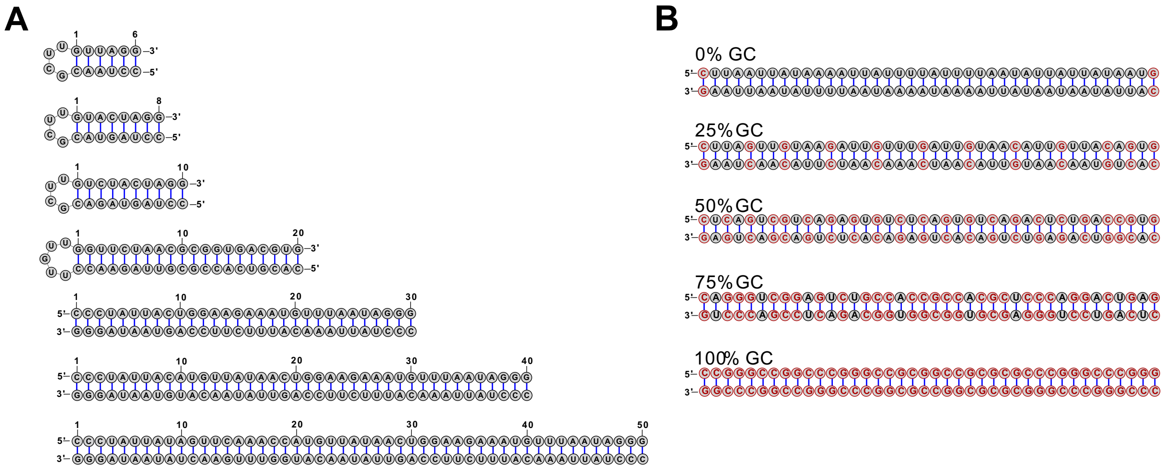


**Figure S1. RNA substrate structure.** (a) RNA substrates with varying length (6bp to 50bp). (b) RNA substrate with varying GC content (0-100%). GC pairs are marked red. All RNAs have 5' fluoresce labels.

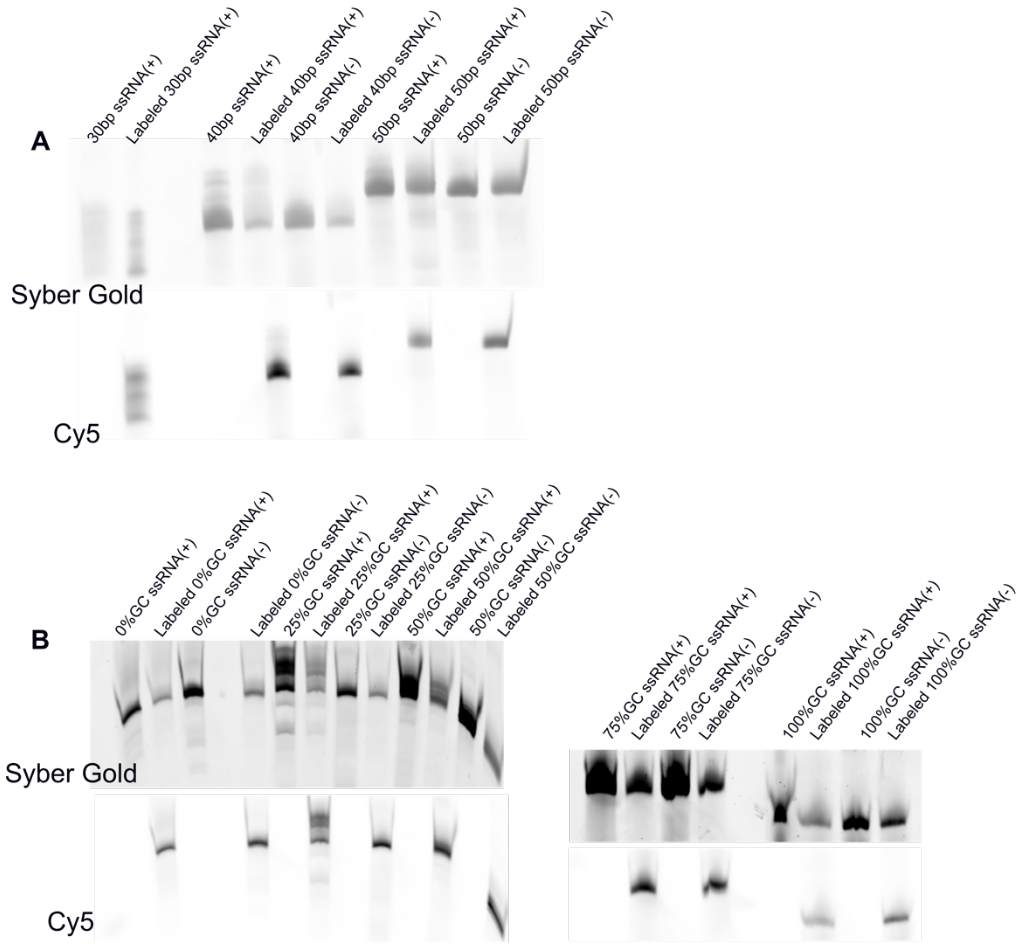


**Figure S2. Validation of RNA substrate labeling by RNA gel electrophoresis.** (a) Gel visualized in Syber Gold showing all RNA substrate with varying length (30bp ssRNA, 40bp ssRNA, and 50bp ssRNA) (*top*). Labeled RNAs with varying length are visualized in Cy5 channel (*bottom*). (b) Gel image of RNA substrate with GC content ranging from 0% to 50% *(left)* and 75% to 100% *(right)* are visualized in both Syber Gold channel and Cy5 channel. Labeled ssRNAs are then annealed with unlabeled complementary pair for MST test.

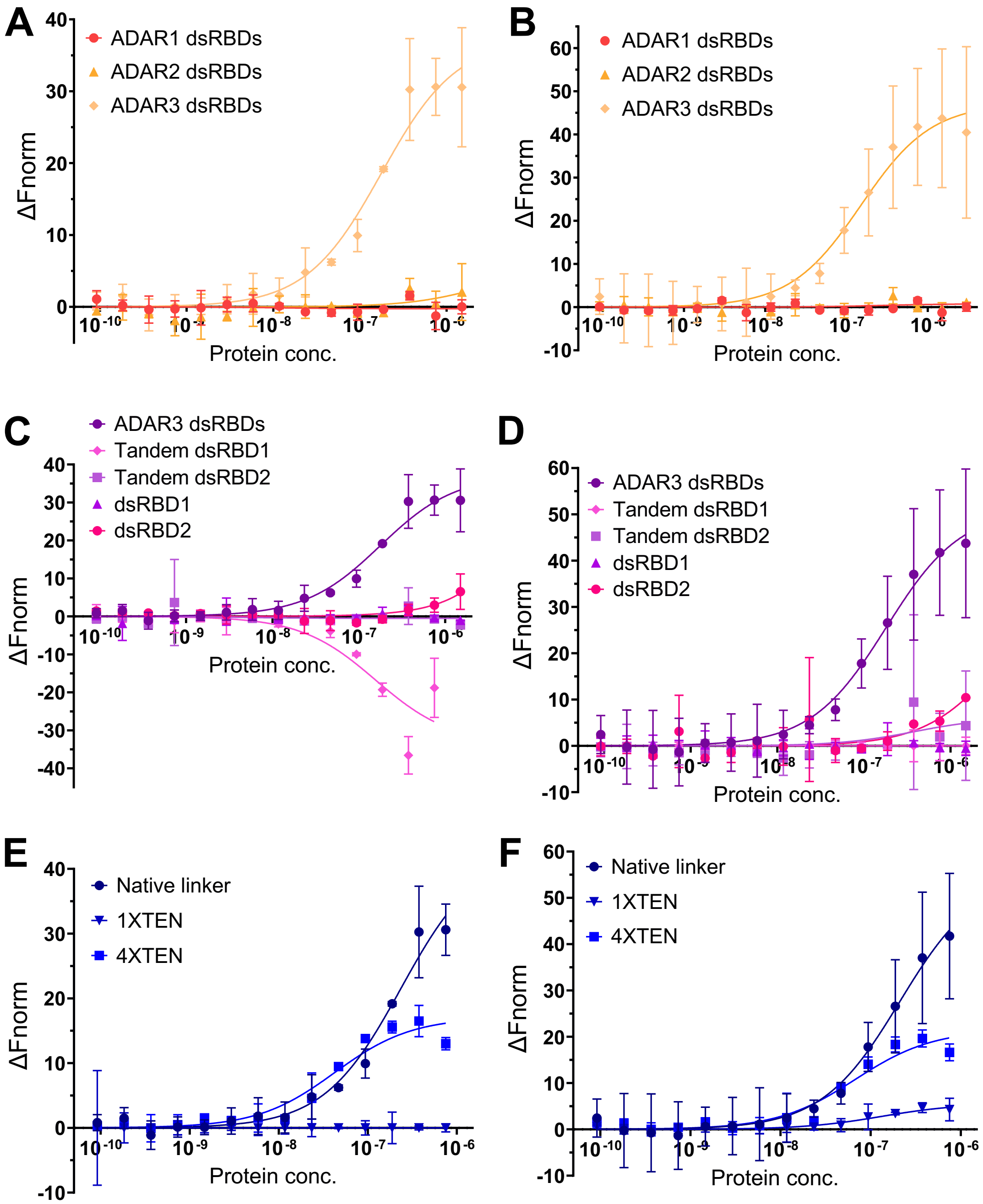


**Figure S3. Characterization of dsRBD binding to random sequence RNAs: MST binding curves showing ΔFnorm responses.** MST binding curves for dsRBDs of ADAR1, ADAR2, and ADAR3 measure with the (a) randomized dsRNA substrate rsRNA-1 and (b) the randomized dsRNA substrate rsRNA-2. MST binding curves for ADAR3 variants, including the wild type dsRBDs, tandem dsRBD1 repeats, tandem dsRBD2 repeats, isolated dsRBD1, and isolated dsRBD2 measured with (c) rsRNA-1 and (d) rsRNA-2. MST binding curves of dsRADAR variants containing different linker length (native, 1x, and 4x) measured with (e) rsRNA-1 and (f) rsRNA-

2. All MST experiments were performed in 20 mM Tris, pH=7.6, 125 mM NaCl, 30 mM KCl, 5 mM MgCl2, 0.5 mM DTT.

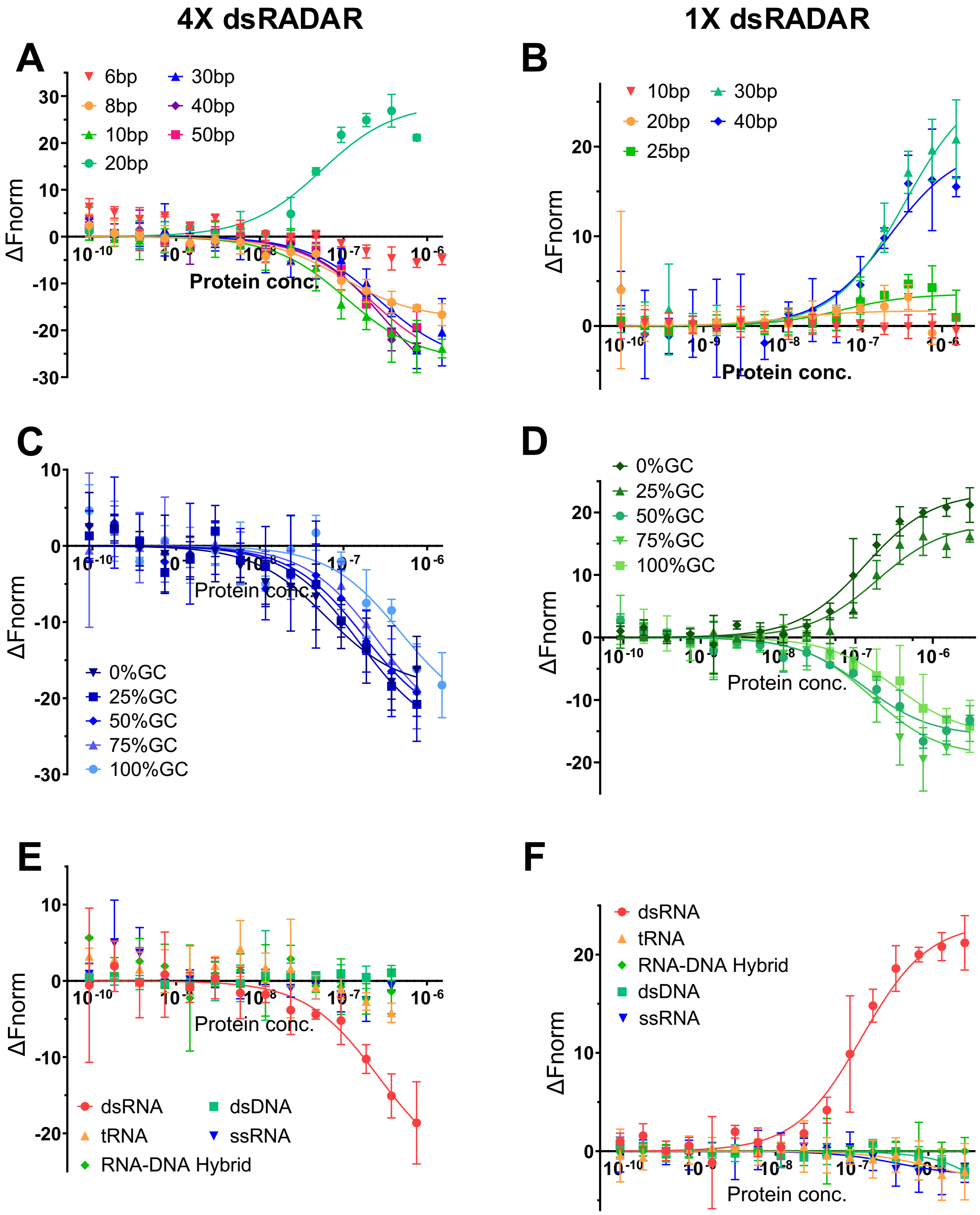


**Figure S4. In vitro characterization of dsRADARs: MST binding curves showing ΔFnorm Responses.** MST binding curves of RNAs having varying duplex length with (a) 4x dsRADAR and (b) 1x dsRADAR. MST binding curves of 40bp dsRNA having varying GC-content with (c) 4x dsRADAR and (d)

1x dsRADAR. MST binding curves of off-target substrates tRNA, RNA-DNA hybrid, dsDNA and ssRNA with (e) 4x dsRADAR and (f) 1x dsRADAR. All MST experiments were performed in 20 mM Tris, pH=7.6, 125 mM NaCl, 30 mM KCl, 5 mM $MgCl_2$, 0.5 mM DTT.

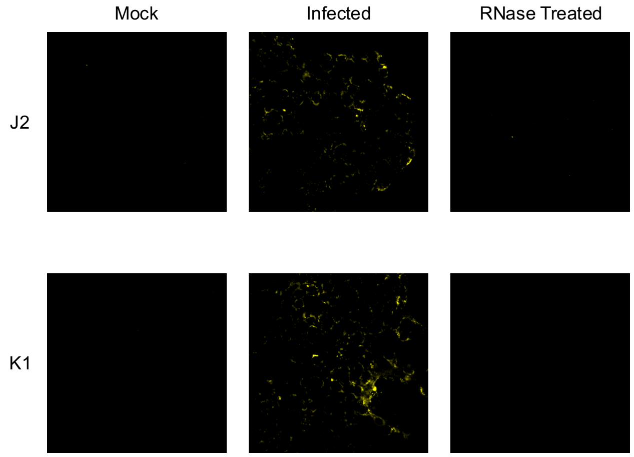


**Figure S5. Immunofluorescence analysis of infected cells after RNase treatment stained with J2 and K1.** Immunofluorescence images of infected cells after RNase treatment (*right*) compared with untreated infected cells (*middle*) and mock cells (*left*).

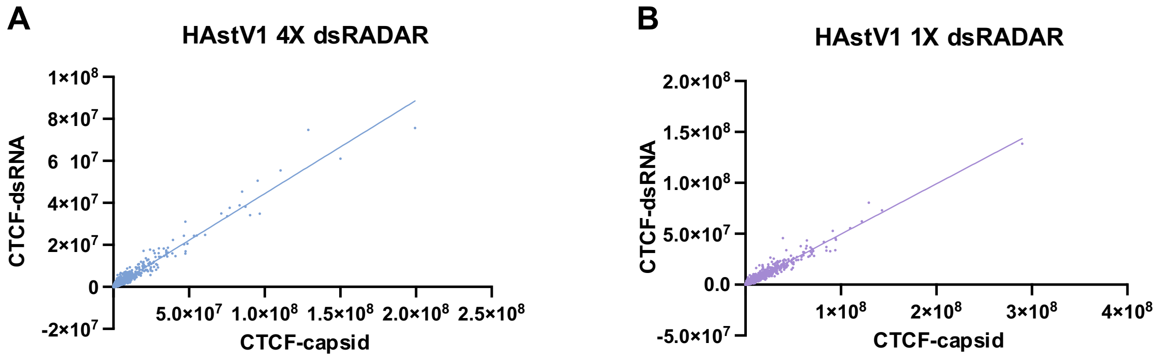


**Figure S6. Linear regression analysis of viral capsid signal and dsRADAR staining.** (a) Simple linear regression calculated based on single cell CTCF of viral capsid and 1x dsRADAR ($p<0.0001$) and (b) 4x dsRADAR ($p<0.0001$).

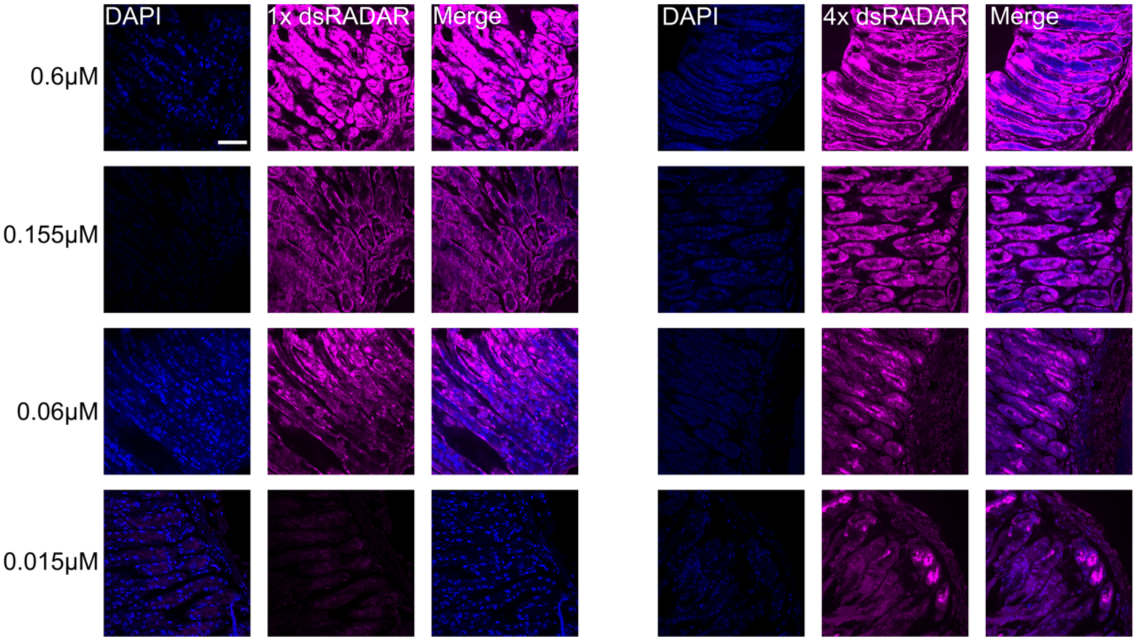


**Figure S7. Optimization of dsRADAR concentration for tissue staining.** FFPE tissue slides from Adar$^{\Delta PC}$, Mavs$^{-/-}$ mouse gastric tissue are used for dsRADAR workflow optimization. Four different concentrations of dsRADAR were tested (0.6μM, 0.155μM, 0.06μM, 0.0155μM). 0.0155μM of dsRADAR was finalized for following studies.
